# Membrane mechanics dictate axonal morphology and function

**DOI:** 10.1101/2023.07.20.549958

**Authors:** Jacqueline M. Griswold, Mayte Bonilla-Quintana, Renee Pepper, Christopher T. Lee, Sumana Raychaudhuri, Siyi Ma, Quan Gan, Sarah Syed, Cuncheng Zhu, Miriam Bell, Mitsuo Suga, Yuuki Yamaguchi, Ronan Chéreau, U. Valentin Nägerl, Graham Knott, Padmini Rangamani, Shigeki Watanabe

## Abstract

Axons are thought to be ultrathin membrane cables of a relatively uniform diameter, designed to conduct electrical signals, or action potentials. Here, we demonstrate that unmyelinated axons are not simple cylindrical tubes. Rather, axons have nanoscopic boutons repeatedly along their length interspersed with a thin cable with a diameter of ∼60 nm like pearls-on-a-string. These boutons are only ∼200 nm in diameter and do not have synaptic contacts or a cluster of synaptic vesicles, hence non-synaptic. Our *in silico* modeling suggests that axon pearling can be explained by the mechanical properties of the membrane including the bending modulus and tension. Consistent with modeling predictions, treatments that disrupt these parameters like hyper- or hypo-tonic solutions, cholesterol removal, and non-muscle myosin II inhibition all alter the degree of axon pearling, suggesting that axon morphology is indeed determined by the membrane mechanics. Intriguingly, neuronal activity modulates the cholesterol level of plasma membrane, leading to shrinkage of axon pearls. Consequently, the conduction velocity of action potentials becomes slower. These data reveal that biophysical forces dictate axon morphology and function and that modulation of membrane mechanics likely underlies plasticity of unmyelinated axons.

## Main Text

Axons are ultrathin membrane tubes designed for rapid conduction of electrical signals, or action potentials (APs), across a tissue or whole organism to convey information. It is well appreciated that AP propagation is impacted by the intricate morphology^1–4^ and cable-like property of axons^5, 6^. During AP conduction in large diameter axons, the diameter of axons increases to lower the axial electrical resistance^4, 7–10^. In the mammalian central nervous system, high-frequency electrical stimulation of axons induced nanoscale remodeling of axonal morphology, where a transient enlargement of synaptic varicosities (by 20%) was followed by a sustained widening of the axons (by 5%)^11^. This, in turn, leads to bidirectional changes in AP conduction velocity, as predicted by cable theory considering the biophysical effects of membrane capacitance and axial resistance. Thus, seemingly minute changes in axon morphology can sensitively tune AP conduction and overall neuronal function.

The morphology of thin membrane tubes like axons is typically dictated by the biophysical properties of the membrane. Electron microscopy (EM) reconstruction of unmyelinated rat hippocampal axons shows that their diameter on average is only 170 ± 40 nm^12^. Such thin membrane tubes should spontaneously form a pearls-on-a-string morphology due to pearling instability, like *in vitro* membrane tubes of the similar diameter^13–20^. Membrane tube pearling is regulated by lipid composition, membrane tension, membrane-protein interactions, and other membrane properties^21^. In fact, pearling behavior is noted in large (2.5 µm diameter) unmyelinated axons when factors such as osmotic pressure^19^ and tension^22^ are modulated. Likewise, unmyelinated axons also exhibit extreme pearling as they degenerate, presumably due to the increase in membrane tension and the loss of cytoskeleton integrity^23^. However, most unmyelinated axons appear relatively uniform in diameter by light microscopy and electron microscopy experiments where the tissue has been fixed^12^. Thus, it is unclear whether small diameter unmyelinated axons in the mammalian central nervous system are uniform in diameter or spontaneously pearl like *in vitro* membrane experiments.

Here we used high-pressure freezing electron microscopy and *in silico* modeling to determine the ultrastructure of axon morphology in mouse neurons. High-pressure freezing is key to studying axon ultrastructure because it circumvents fixation artifacts like membrane distortion and protein aggregation and preserves membrane morphology in a near-native state^24–26^. We find that axons are not simple cylindrical tubes, but rather exhibit nanoscopic pearls-on-a-string morphology. Ultrathin axon tracts of ∼60 nm connect the pearled regions, or non-synaptic boutons (NSBs), of ∼200 nm. *In silico* modeling shows that pearling minimizes bending energy and further predicts that pearled morphology arises from biophysical properties governing membrane mechanics. Indeed, experimental manipulation of osmotic pressure, the cytoskeleton, and membrane fluidity all alter pearling behavior of axons, indicating that axon pearling is parameterized by various biophysical factors. Based on this morphology, we modeled the action potential propagation using the cable equation with Hodgkin and Huxley currents. We found that manipulation of biophysical factors changes the conduction velocity of action potentials. Thus, our study provides deeper insight into how axon morphology and function are controlled by a delicate balance of biophysical forces acting on the plasma membrane.

## Results

### Axons are pearled, not tubular, under physiological conditions

Membrane tubes *in vitro* typically form pearling due to tension driven instability^14^. Although ultrathin membrane tubes like unmyelinated axons are susceptible to such biophysical changes, previous work suggest that axons are tubular^12, 27–31^. However, ultrastructural analysis is typically performed on samples prepared with aldehyde-based fixatives under conditions that do not preserve fine morphology. Thus, to interrogate the morphology of unmyelinated axons, we performed high-pressure freezing and electron microscopy analysis of neurons from acutely extracted mouse brain tissue (Postnatal days 42-70, P 42-70, as per^32^), organotypic slice culture, and dissociated hippocampal culture from embryonic day 18 (E18) mouse pups. These three types of tissue preparation were chosen since they have different biophysical environments and may display distinct morphologies^33^.

As in previous studies^12, 27, 34^, axons appeared cylindrical in chemically fixed tissues (Extended Data Fig. 1a,b). By contrast, when prepared using the high-pressure freezing method, axons in all preparation types exhibit a pearls-on-a-string morphology (Fig. 1a,b and Extended Data Fig. 1a-d) with changes in dimension depending on culturing conditions. Pearled regions are defined as non-synaptic boutons (NSBs) and the region between two NSBs is designated the connector with boundaries defined by the inflection points (Fig. 1b, c). To quantify this morphology, the width and length of NSBs and connectors were measured. The axonal morphology of acute slices and organotypic slices was similar. NSB dimensions on average in acute slices were 360 ± 10 nm long by 200 ± 6 nm wide and in organotypic slices were 340 ± 7 nm long by 180 ± 3 nm wide (Fig. 1d and Extended Data Fig. 1d). Connector dimensions on average in acute slices were 380 ±15 nm long by 58 ± 1 nm wide and in organotypic slices were 250 ± 10 nm long by 50 ± 2 nm wide (Fig. 1d and Extended Data Fig. 1d). Cultured neurons were slightly larger in all dimensions with the NSBs being 480 ± 9 nm long by 275 ± 5 nm wide and connectors being 410 ± 15 nm long by 110 ± 2 nm wide (Fig. 1d and Extended Data Fig. 1d), most likely reflecting the changes in biophysical properties associated with substrate stiffness^33^. Nonetheless, these results suggest that unmyelinated axons indeed exhibit membrane pearling due to tension-driven instability.

**Fig. 1.**
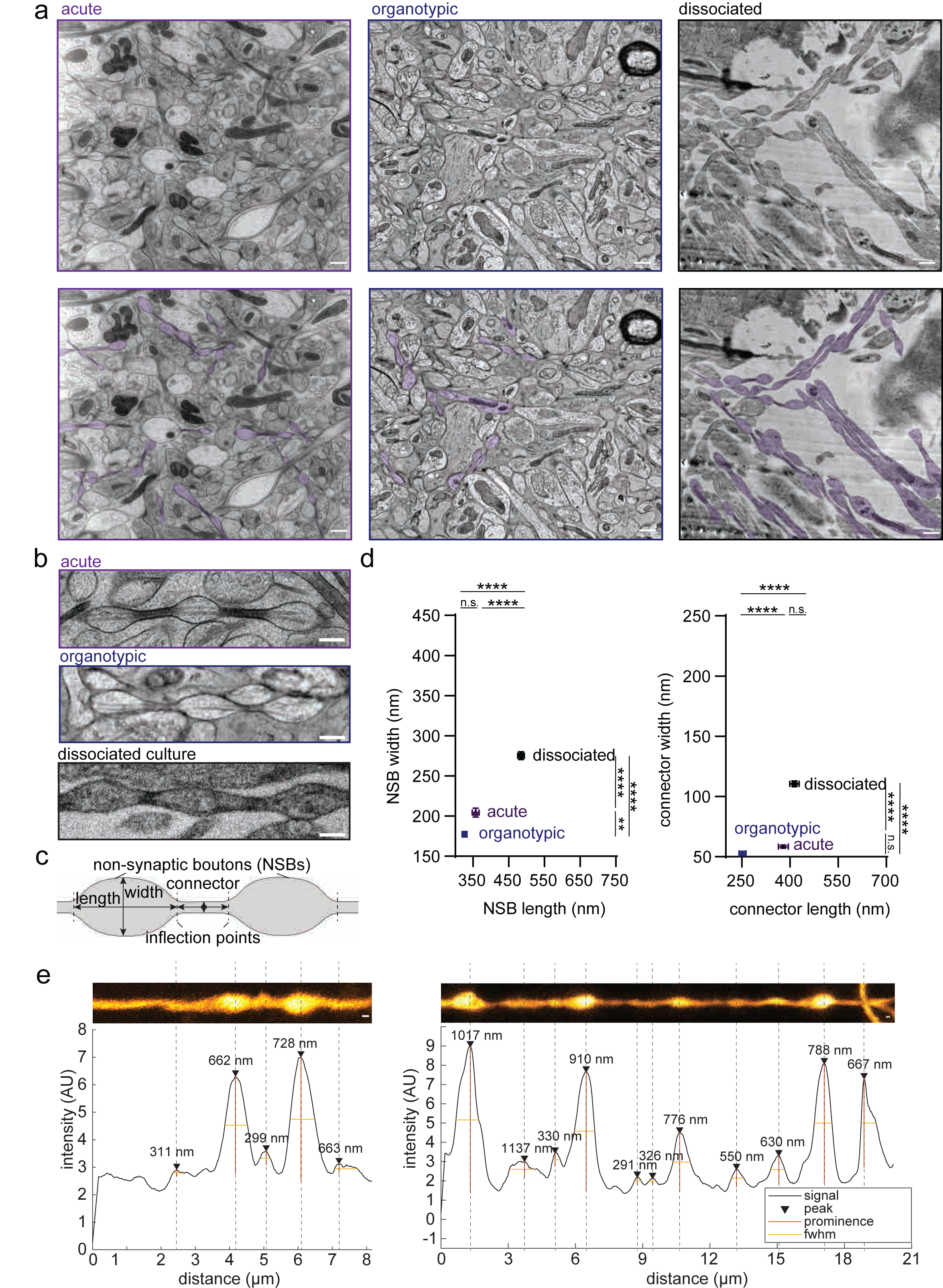
Axons are pearled, not tubular, under homeostatic conditions. a) Representative electron micrographs from acutely extracted mouse brain tissue (left), organotypic slice cultures of mouse hippocampus (middle), and dissociated mouse hippocampal neuronal culture after high pressure freezing (HPF). Some axons in each micrograph are traced and color-coded on the bottom panel. Scale bar: 500 nm. b) High magnification images of axons, representative of each condition. More example micrographs are found in Extended Data Fig. 1a-c. Scale bar: 200 nm. c) A schematic showing two non-synaptic boutons flanked by a connector. Inflection points define the boundary between these two features. Both width and length are measured at NSBs and connectors, as shown. d) Plots showing dimensions of NSBs (left) and connectors (right) from indicated tissue types. Dimensions are measured from three independent samples for acutely extracted brain tissue and dissociated neuron culture and one for organotypic slices. n = 30 axons from each acutely extracted sample, n = 133 axons from the organotypic sample, n = 100 axons from each dissociated sample. Super plots showing variability are available in Extended Data Fig. 1d. Mean and S.E.M. are plotted. Kruskal-Wallis test, followed by Dunn’s multiple comparison test. *n.s.,* not significant*, ** P <0.01, ****P* < 0.0001. e) Representative STED micrographs showing axons from organotypic slice cultures of mouse hippocampi. The numbers represent the length of each NSB, measured at full-width half- maximum using Matlab scripts. Numbers indicate the measured length at each NSB. Scale bar: 200 nm.

To ensure that axon pearling was not induced by our experimental conditions, we performed live-cell superresolution imaging of organotypic mouse hippocampal slice cultures at 21-35 days *in vitro* (DIV 21-35). Cytosolic GFP was expressed using Sindbis virus with stereotaxic injection in the CA3 area of hippocampal slices ∼36 hours prior to the experiments. Stimulated Emission Depletion (STED) microscopy analysis of live tissues show the pearling behavior of axons, similarly to the morphology observed by electron microscopy (Fig. 1e and Extended Data Fig. 1e). These data suggest that nano-pearling is a prominent feature of unmyelinated axons.

### Membrane mechanics regulate axon morphology

We first considered whether the plasma membrane material properties could be driving the experimentally observed nano-pearled shape. Building upon the rich literature of membrane continuum mechanics, we constructed a model for predicting the membrane shapes of unmyelinated axons (Fig. 2a). We model the membrane as a thin elastic surface with energy given by the equation in Fig. 2a (see Table 1 for notation); this is the classic Helfrich Hamiltonian, which represents the elastic energy as a function of the surface curvatures^35, 36^.

**Fig. 2.**
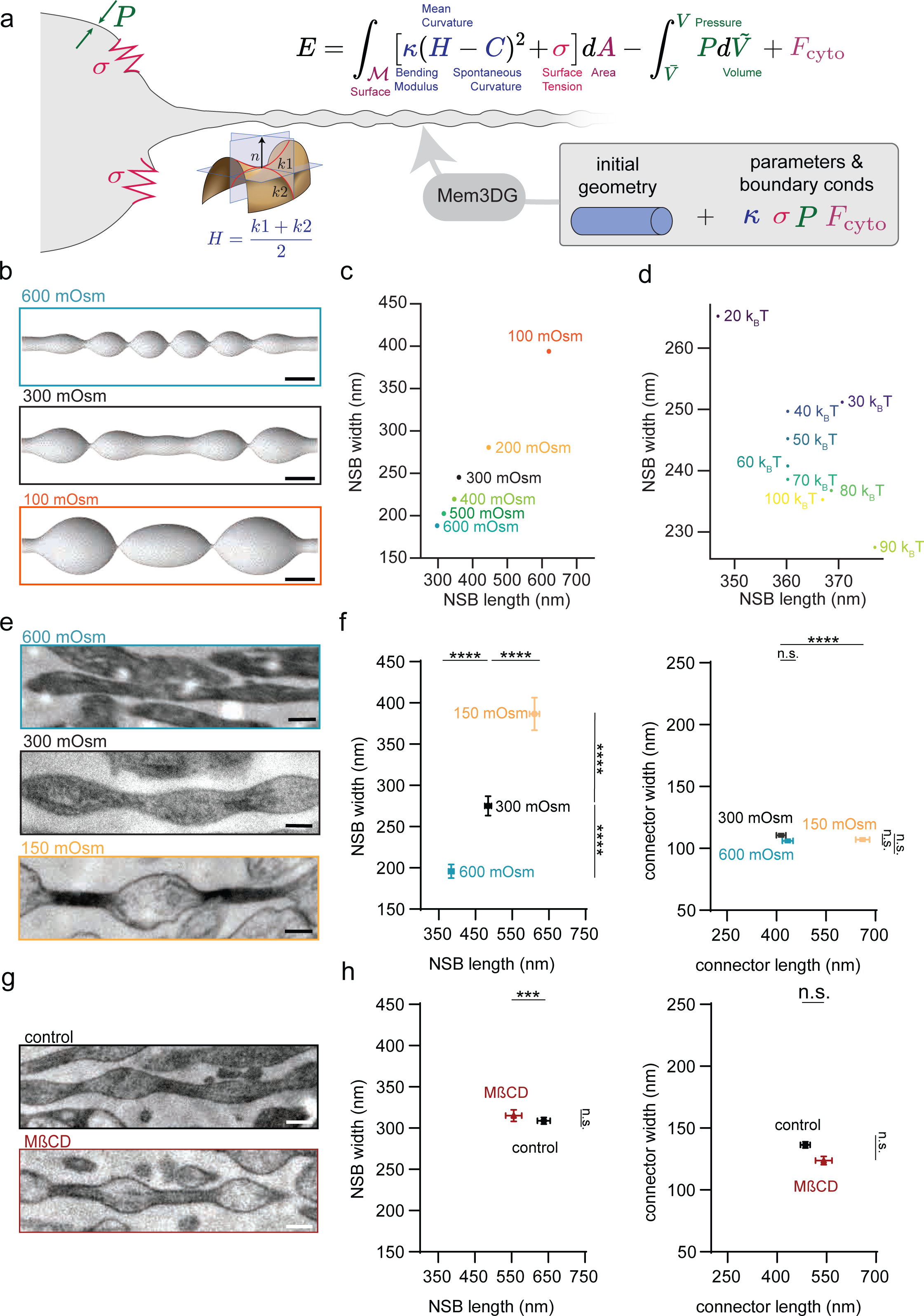
Membrane mechanics dictate axon pearling. a) Axon morphology is modeled using classic Helfrich membrane model and governed by the membrane bending, surface tension, and osmotic condition. b) Model prediction of axon morphology in indicated osmotic conditions. Scale bar: 200 nm. c) Plot showing the dimensions of NSBs at indicated osmotic conditions. Note that the NSB size is inversely scaled with the external osmotic pressure. d) Plot showing the dimensions of NSBs with varying membrane rigidity, ranging from 20 kBT to 100 kBT. e) Example micrographs of axons high-pressure frozen at indicated osmotic conditions. More example micrographs are found in Extended Data Fig. 3a. Scale bar: 200 nm. f) Plots showing the dimensions of NSBs (left) and connectors (right) from neurons in (e). n = 100 axons from 3 replicates. Super plots showing variability are available in Extended Data Fig. 3b. Mean and S.E.M. are plotted. Kruskal-Wallis test, followed by Dunn’s multiple comparison test. *n.s.,* not significant*, ****P* < 0.0001. g) Example micrographs of axons from cultured neurons treated with sham (control) or 5 mM MßCD for 30 min. More example micrographs are available in Extended Data Fig. 4c. Scale bar: 200 nm. h) Plots showing the dimensions of NSBs (left) and connectors (right) from neurons in (g). n = 100 axons from 3 replicates. Super plots showing variability are available in Extended Data Fig. 4d. Mean and S.E.M. are plotted. Kruskal-Wallis test, followed by Dunn’s multiple comparison test. *n.s.,* not significant*, ***P* < 0.001.

Minimizing the energy with respect to the shape yields solutions for mechanical equilibrium^37–39^. We used a computational approach, Mem3DG^40^, which represents the membrane using triangulated meshes and calculates the Helfrich energy and corresponding forces using strategies from discrete differential geometry.

Starting from an open-ended cylindrical tube connected to implicit membrane reservoirs, we vary the osmotic pressure, bending rigidity, and membrane tension systematically and obtain relaxed geometries (Extended Data Fig. 2). Geometries for all conditions exhibited nano-pearled morphologies with varying varicosity. Consistent with prior work on membrane tube mechanics^14, 15, 19, 21, 41, 42^, manipulation of osmotic pressure on membranes has a strong influence on nano-pearled morphology. Increasing solvent osmolarity reduces NSB width and length (Fig. 2b, c). Raising tension also constrains the ability of the system to increase surface area leading to reduced NSB width and length (Extended Data Fig. 2c,d), albeit at a rate subleading the osmotic pressure contribution (Extended Data Fig. 2). The bending rigidity on the other hand had a modest effect on NSB geometry (Fig. 2d). With the assumption that model parameters such as spontaneous curvature, tension, and bending rigidity are homogeneous, the output geometries are limited to periodic unduloid-like shapes^16, 20^, and thus, we cannot capture the behavior of connectors, which would require the introduction of arbitrary heterogeneity^21^ or additional speculative physics. In summary, our simulations predict that NSBs driven by a membrane pearling instability scale inversely with osmotic condition: increasing the osmolarity of the milieu decreases both the NSB length and width.

To test modeling predictions, ultrastructural analysis was performed on dissociated hippocampal culture (DIV 21), high-pressure frozen while manipulating the osmotic pressure from isotonic conditions to either hyper- (600 mOsm) or hypo- tonic (150 mOsm) conditions (Fig. 2d,e). In general, hyperosmotic solution would shrink membranes, resulting in reduction in membrane tension, while hypoosmotic solution would do the opposite. As in modeling (Fig. 2b,c), the dimensions of NSBs were inversely correlated with the osmolarity (Fig. 2e,f and Extended Data Fig. 3a,b). Upon doubling the osmolarity the NSBs shrunk by 45% in width (275 ± 5 nm to 190 ± 4 nm wide) and by 25% in length (480 ± 9 nm to 370 ± 8 nm long) (Fig. 2e,f and Extended Data Fig. 3a,b). Upon halving the osmolarity there was an increase in NSB width of 45% (275 ± 5 nm to 390 ± 8 nm wide) and length by 35% (480 ± 9 nm to 610 ± 13 nm long) (Fig. 2e,f and Extended Data Fig. 3a,b). In hyperosmotic conditions, the dimensions of connectors did not change (410 ± 15 nm to 430 ± 16 nm long and 111 ± 2 nm to 106 ± 3 nm wide), while in hypoosmotic conditions, the connector length increased by 53% (410 ± 15 nm to 661 ± 21 nm long) but not width (111 ± 2 nm to 107 ± 3 nm wide). Nevertheless, the experimental observation that NSB width and length correlate inversely with solvent osmolarity is in accord with the membrane mechanics model, suggesting that membrane pearling instability may underlie the experimentally observed pearled shapes.

**Fig. 3.**
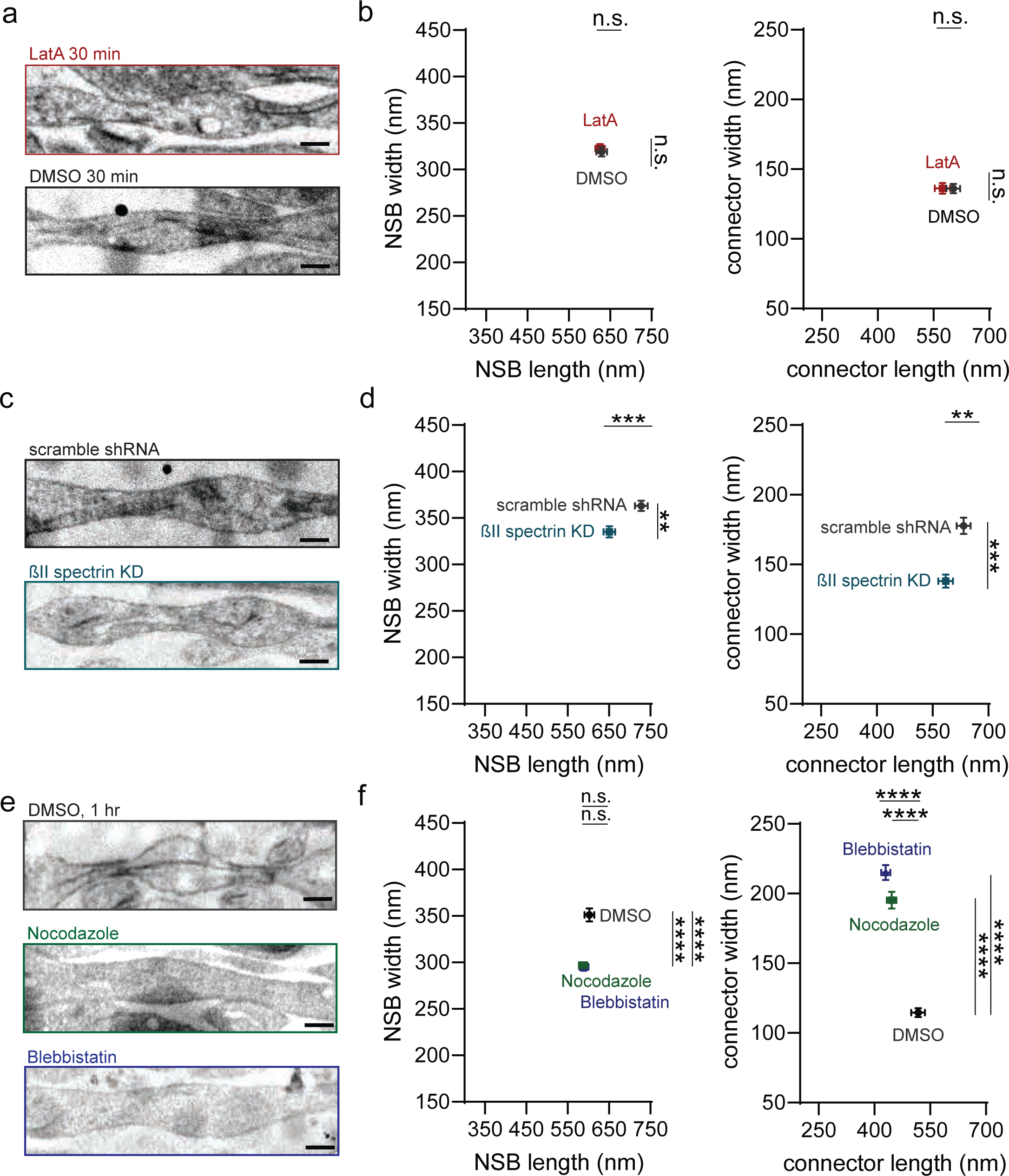
MPS is not sufficient to explain pearled axon morphology. a) Example micrographs of axons from cultured mouse hippocampal neurons treated with either 0.1% DMSO or 20 µM Latrunculin A (LatA) for 30 min. Scale bar: 200 nm. b) Plots showing dimensions of NSBs (left) and connectors (right) from axons in (a). DMSO: NSB length 630 ± 13 nm, NSB width 320 ± 5 nm, conn. length 600 ± 18 nm, conn. width 136 ± 3 nm. LatA: NSB length 230 ± 11 nm, NSB width 320 ± 5 nm, conn. length 570 ± 20 nm, conn. width 136 ± 4 nm. c) Example micrographs of axons from neurons infected with lentivirus carrying either scramble or ßII Spectrin (Sptbn1) shRNA. Scale bar: 200 nm. d) Plots showing dimensions of NSBs (left) and connectors (right) from axons in (c). e) Example micrographs of axons from neurons treated with 0.1% DMSO, 50 µM Nocodazole, or 10 µM Blebbistatin for 1 hr. Scale bar: 200 nm. f) Plots showing dimensions of NSBs (left) and connectors (right) from axons in (e). DMSO: NSB length 600 ± 12 nm, NSB width 350 ± 7 nm, conn. length 520 ± 19 nm, conn. width 115 ± 3 nm. Nocodazole: NSB length 590 ± 10 nm, NSB width 300 ± 3 nm, conn. length 450 ± 13 nm, conn. width 200 ± 6 nm. Blebbistatin: NSB length 590 ± 11, NSB width 290 ± 4 nm, conn. length 430 ± 12 nm, conn. width 215 ± 5 nm. In each experiment, N = 3 independent cultures and n = 300 axons. Super plots showing variability are available in Extended Data Fig. 3. Mean and S.E.M. are plotted. All conditions in Fig. 3 are analyzed at the same time, and thus, Kruskal-Wallis test, followed by Dunn’s multiple comparison test was used. *n.s.*, not significant, \*\**P* < 0.01,\*\*\**P* < 0.001, \*\*\*\**P* < 0.0001.

Since these osmolarity changes are extreme, the experiments were repeated using more physiologically relevant osmolarity changes (280 mOsm to 400 mOsm) by replacing glucose with the membrane impermeable solute mannitol and varying its concentration (see Methods for detail). As expected, the effects were less pronounced, but trends were similar: the hypertonicity causes NSBs to shrink in length by 5% (510 ± 10 nm to 470 ± 8 nm) but not in width (250 ± 4 nm to 230 ± 3 nm), while decreasing the osmolarity caused the NSBs to expand (width: increase by 20%, length: increase by 12%) (Extended Data Fig. 3c-e). Connector dimensions did not change in hypertonic conditions (490 ± 19 nm to 560 ± 13 nm long and 114 ± 3 nm to 118 ± 3 nm wide) or in hypotonic conditions (490 ± 19 nm to 540 ±16 long and 114 ± 3 nm to 125 ± 3 nm wide) (Extended Data Fig. 3c-e). Therefore, osmotic pressure and thereby membrane tension, in part, regulate pearled axon morphology.

To further test the contribution of membrane mechanics, we manipulated membrane fluidity using methyl-ꞵ-cyclodextrin (MꞵCD, 5 mM, 30 min) to remove the cholesterol from the plasma membrane of DIV 21 cultured hippocampal neurons^43, 44^. Assuming that cholesterol imparts a general stiffening of the membrane, the removal of cholesterol by MβCD treatment would produce a decrease in membrane bending rigidity, which in turn, produces a general decrease in NSB size based on modeling (Fig. 2d). Upon MꞵCD treatment, the cholesterol level on axons was probed by the exogenously applied ALOD4-NeonGreen^45^. Within 30 min, essentially all accessible cholesterol was removed from the plasma membrane (Extended Fig. 4a,b). Electron microscopy analysis showed that the NSB length decreases from 640 ± 17 nm in the control conditions to 560 ± 14 nm (12.5% decrease; Fig. 2g,h and Extended Data Fig. 4c,d), indicating that membrane fluidity and rigidity also contributes to pearled axon morphology. The remaining dimensions did not change (NSB width: 310 ± 7 nm to 310 ± 8 nm, conn. length: 490 ± 15 nm to 540 ± 20 nm) (Fig. 2g,h and Extended Data Fig. 4c,d). The concurrence of axon geometry with respect to experimental and model perturbations suggests that membrane mechanics may be a key driver of axon nano-pearling.

**Fig. 4.**
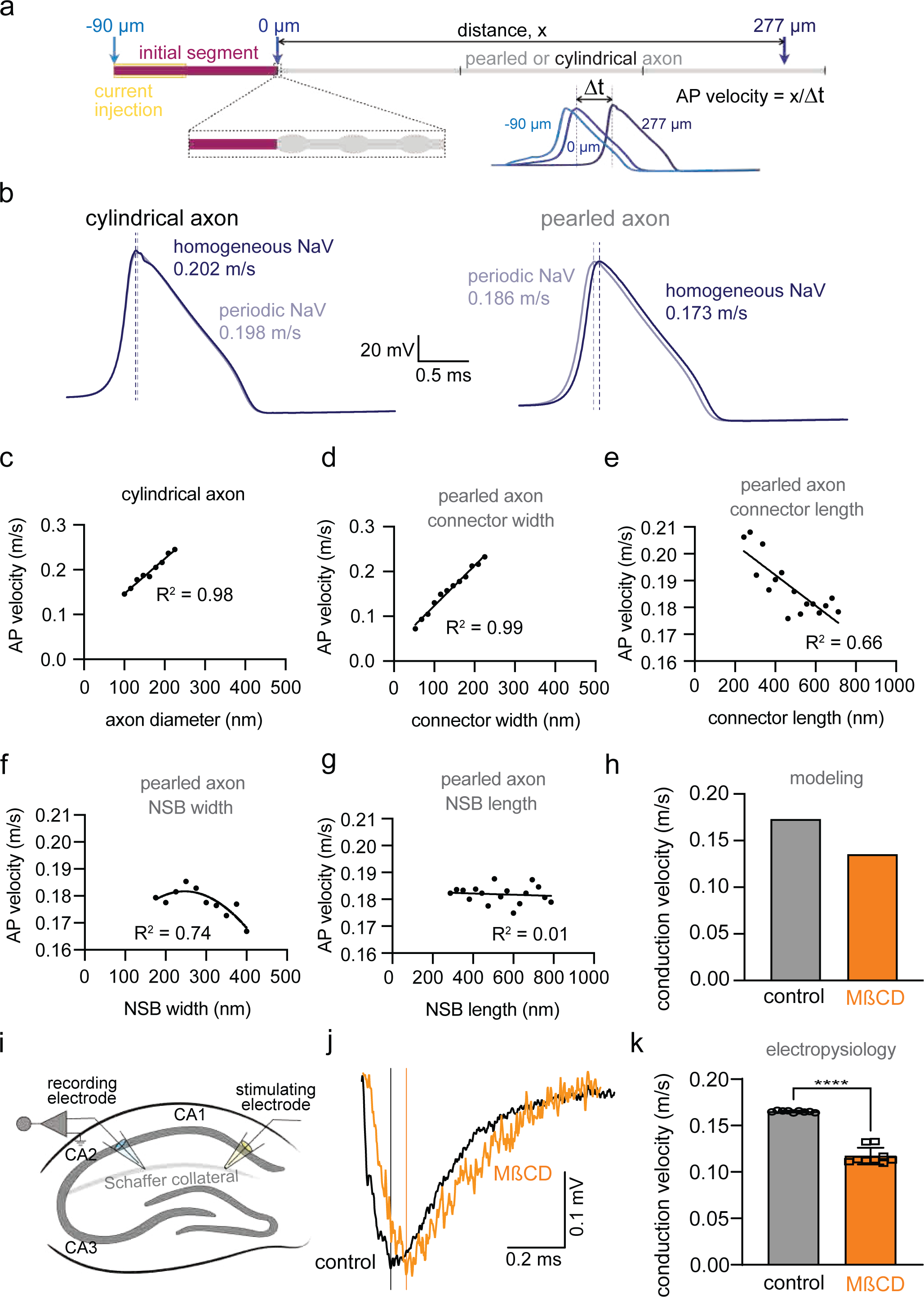
Action potential propagation relies on axonal morphology. a. Schematic showing the model setup. Action potentials are modeled in real geometries using a generalized cable equation to calculate the spatial and temporal distribution of channel current, membrane voltage, and the gating variables. An axon initial segment (AIS) of 90 µm is connected to either a cylindrical or a pearled axon of 300 µm. Currents are injected at the first half of the AIS for 1 ms to induce an action potential. The resulting voltage change is measured at -90 µm, 0 µm, and +270 µm points relative to the end of the AIS. The velocity is determined from the time it took from the peak of the voltage response at 0 µm to another peak at 270 µm. b. Voltage responses at 270 µm from cylindrical axons (left) and pearled axons (right), when voltage-gated sodium channels (NaV) are distributed either uniformly (dark color) or periodically (lighter color). Note that the distribution of NaV only matters if axons are pearled. c. Plot showing the relationship of action potential (AP) velocity to the diameter of cylindrical axons. Dots are fitted with a simple linear regression curve. d-g. Plots showing the relationship of action potential (AP) velocity to the connector width (d), connector length (e), NSB width (f), and NSB length (g). Dots are fitted with a simple linear regression curve, except for (f), which is fitted by a non-linear gaussian curve. h. Plot showing the predicted AP conduction velocity based on the dimensions of NSBs and connectors in neurons treated with sham (control) or 5 mM MßCD. i. Schematic showing the electrophysiology recording set up. The Schaffer Collateral was stimulated from the end of CA1 to measure the back-propagating action potential in the CA1 near the CA2. j. Example traces from the recording in acute slices of mouse hippocampus, treated with either sham (control) or 5 mM MßCD for 30 min. A solid vertical line is placed at the peak. k. Plot showing the AP conduction velocity from the experiments in (j). Mann-Whitney U test. N = 4 animals each, n = 4 slices. \*\*\*\**P* < 0.0001.

### Non-muscle myosin II contractility contributes to pearled axon morphology

Although the aforementioned model is inspired by membrane mechanics, parameters such as the effective membrane tension may represent other mechanical contributions.

Membrane tension is a sum of in-plane tension and cortical cytoskeleton attachment^46–48^. In axons, actin forms periodic rings, termed membrane periodic cytoskeleton (MPS), by its interaction with spectrin^49–51^. To discern the contribution of the MPS, we tested the role of different cytoskeletal components. Ultrastructural analysis of mouse hippocampal neurons (DIV 21) was performed after treatment with DMSO (0.1%, 30 min) or Latrunculin A (LatA, 50 µM, 30 min)^49^, which blocks actin dynamics^52^. Expression of ßII spectrin was knocked down with shRNA using lentivirus, as previously described (Extended Data Fig. 5a)^49^. Scramble (scr) shRNA was used as a control. These treatments are shown to perturb the MPS^49^. However, axon nano- pearling was not altered in neurons treated with LatA (Fig. 3a,b and Extended Data Fig. 5b-e).

**Fig. 5.**
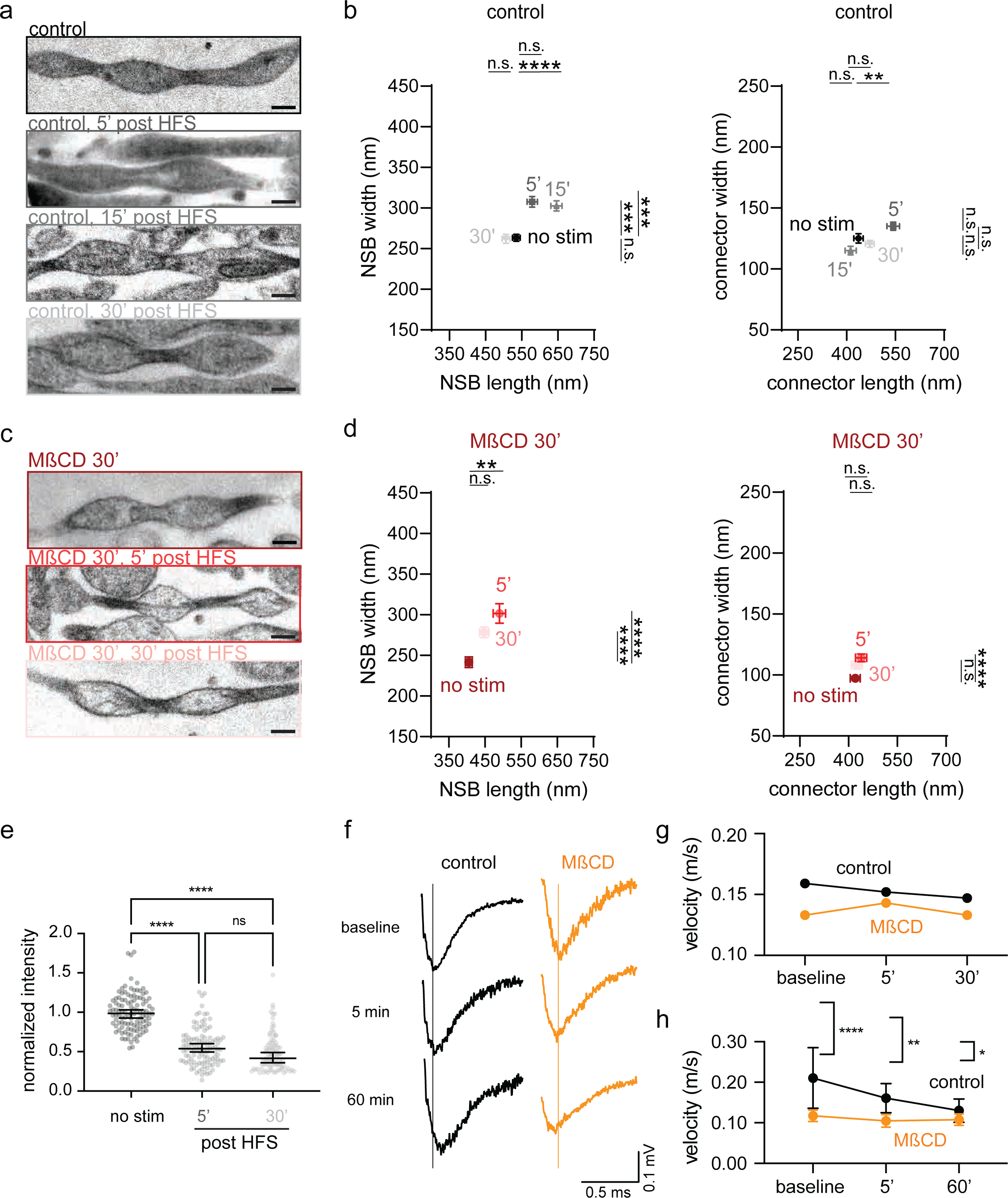
Axonal plasticity is mediated by modulation of membrane mechanics. a. Example micrographs showing axon morphology from control neurons unstimulated or stimulated with 3x100 pulses at 100 Hz (high-frequency stimulation, HFS) and high-pressure frozen at 5 min and 30 min after stimulation. Scale bar: 200 nm. b. Plots showing dimensions of NSBs (left) and connectors (right) from axons in (a). N = 3 independent cultures except for control 15’ post stim which was N = 2. n = 100 axons each. *n.s.,* not significant, \*\**P* < 0.01, *\*\*\*\*P* < 0.0001. c. Example micrographs showing axon morphology from MßCD-treated neurons (5 mM, 30 min) unstimulated or stimulated with 3x100 pulses at 100 Hz (high-frequency stimulation, HFS) and high-pressure frozen at 5 min and 30 min after stimulation. Scale bar: 200 nm. d. Plots showing dimensions of NSBs (left) and connectors (right) from axons in (d). no stim: NSB length 400 ± 10 nm, NSB width 240 ± 7 nm, conn. length 420 ± 15 nm, conn. width 97 ± 3 nm; post HFS 5’: NSB length 490 ± 17 nm, NSB width 300 ± 12 nm, conn. length 440 ± 17 nm, conn. width 114 ± 3 nm; post HFS 30’: NSB length 450 ± 13 nm, NSB width 280 ± 7 nm, conn. length 430 ± 16 nm, conn. width 108 ± 3 nm. N = 3 independent cultures except for MβCD 5’ post stim which was N = 2. n = 100 axons each. *n.s.,* not significant, \*\**P* < 0.01, *\*\*\*\*P* < 0.0001. e. Plots showing the normalized intensity of the cholesterol biosensor NeonGreen-ALOD4 in neurons unstimulated or stimulated with HFS and fixed 5 or 30 min after. Median and 95% confidence interval shown (no stim: 0.99, 95%CI 0.95, 1.04, 5’ post HFS: 0.54 95%CI 0.52, 0.61, 30’ post HFS: 0.42, 95%CI 0.44, 0.54). N = 3 independent cultures. n = >91 axons total. Kruskal-Wallis test, followed by Dunn’s multiple comparison test. *n.s.,* not significant*, ****P* < 0.0001. f. Example traces from electrophysiology experiments in acute slices of mouse hippocampus, performed as described in Fig. 4i. Following baseline measurement, either sham (control) or MßCD was administered for 30 min. Additional recordings were performed after HFS at 5 and 50 min after the initial administration, but the sham or MßCD were applied throughout these additional recordings. g. The predicted AP conduction velocity based on the dimensions of NSBs and connectors before, 5 min after, and 30 min after HFS. h. The measured AP conduction velocity, before, 5 min after, and 30 min after HFS. N = 4 animals each, n = 4 slices. Mann-Whitney U test. N = 4 animals each, n = 4 slices. \**P* < 0.05, \*\**P* < 0.01, *\*\*\*\*P* < 0.0001.

Likewise, the pearled morphology remains in ßII spectrin KD neurons (Fig. 3c,d and Extended Data Fig. 5f,g), although axons appeared to shrink when compared to those from the scramble shRNA control: NSB length decreased by 10% (728 ± 15 nm to 650 ± 14 nm, respectively), while the width decreased by 8% (368 ± 6 nm to 335 ± 6 nm, respectively). The connector width also decreased by 22% (178 ± 6 nm to 138 ± 5 nm, respectively) while connector length did not change (630 ± 19 nm to 590 ± 19 nm, respectively). However, the resulting dimensions of axons in ßII spectrin KD neurons are similar to those found in wild-type neurons, suggesting that lentiviral infection may have caused enlargement of axons. However, the overall axon nano-pearling remained after these treatments, suggesting that the MPS plays a minimal role in determining axon morphology.

Given that non-muscle myosin II (NMII) imparts a contractive effect on the axon to potentially induce nano-pearling, we next tested the role of NMII by electron microscopy. We used NMII inhibitor Blebbistatin (10 µM, 1 hr) to block NMII activity^51^. After this treatment, nano- pearling became less pronounced, particularly at the connectors (NSBs width decreased by 23%, connectors width increased by 68%; Fig. 3e,f and Extended Data Fig. 5h,i), suggesting that force generated by NMII contractility likely determines the lower bounds of axon morphology.

Actin-myosin complexes (actomyosin) interplay with microtubules to control cellular architecture^51^. Previous studies show that disruption of microtubules causes dissociation of MPS^50^, suggesting a close link between actomyosin and microtubules within axons. Thus, the role of microtubules in axon morphology was tested by inhibiting microtubule polymerization with Nocodazole (10 µM, 1 hr). Like in NMII inhibition, this treatment had little effect on NSBs but decreased the connector length by 20% while increasing the width by 58% (Fig. 3e,f and Extended Data Fig. 5h,i). Together, these results suggest that the MPS is not required for axon nano-pearling, but generation of non-uniform force due to the periodic organization of the MPS may determine the connector morphology.

### Axon pearling increases the dynamic range of action potential conduction velocity

To determine how nano-pearled axon morphology influences function, we implemented the generalized cable equation with Hodgkin and Huxley currents^1, 53^, taking the pearled morphology into account. We simulated electrical conductance over 300 µm of either pearled or cylindrical axon while injecting currents of 30-40 µA/cm^2^ at the first half of an axon initial segment (AIS, Fig. 4a) and measuring the resulting voltage change at the tip of the AIS (-90 µm), at the end of the AIS (0 µm), and near the end of the model axon (277 µm). The conduction velocity was determined based on the time it took for the voltage change to reach its peak at 0 µm and 277 µm (Fig. 4a). Since voltage-gated sodium channels (NaV) are organized by spectrin and Ankyrin, which make up the MPS, we also simulated APs with NaV placed either uniformly along the axon or periodically with the interspacing of 190 nm^49^. For cylindrical axons, the periodic or uniform distribution of NaVs did not alter the AP conduction velocity (Fig. 4b). However, in nano-pearled axons with the average geometries measured in our experiments, the AP is faster if NaVs are periodically distributed by the MPS.

For cylindrical axons, the larger the diameter, the faster the AP conduction velocity (Fig. 4b,c: the AP velocity = 0.202 m/s). By contrast, the conduction velocity was highly variable in nano-pearled axons, depending on the dimensions of NSBs and connectors. Like in cylindrical axons, there is a linear relationship between the AP velocity and the connector width (Fig. 4d) likely due to the change in axial resistance. Similarly, the connector length has an inverse linear relationship with AP velocity (Fig. 4e). The relationship between NSB width and AP velocity can be described by a concave function with the AP velocity increasing up to the width of 250 nm, after which it decreases (Fig. 4f). However, no clear correlation is observed between NSB length alone and AP velocity, but the ratio between NSB length and width is a key determinant of AP velocity (Extended Data Fig. 6). Thus, based on the local biophysical environment and dimensions of individual NSBs and connectors, the AP velocity can be highly modulable.

To validate our model, we used acute hippocampal slices from P30-40 mouse and recorded fiber volley conduction velocity before and after MꞵCD treatment (5 µM, 30 min). Based on the dimensions of NSBs and connectors in sham (control) and MꞵCD treated neurons (Fig. 2g,h), our model predicts APs to be slower in MꞵCD treated axons (0.135 m/s, ∼22% decrease, NSB 573 x 310 nm, conn. 567 x 121 nm) compared to untreated axons (0.173 m/s, NSB 638 x 309 nm, conn. 486 x 136 nm; Fig. 4h). As predicted by the model, the action potential velocity decreased from 0.164 ± 0.3 m/s to 0.117 ± 3 m/s upon MꞵCD treatment (∼28% decrease; Fig. 4i-k), suggesting a tight correlation between axon nano-pearling and function.

### Axonal plasticity is induced by modulation of biophysical factors

Axon morphology is tightly coupled to neuronal activity and can modulate AP conduction velocity. To better understand how this pearled morphology behaves during sustained neuronal activity we applied three trains of high-frequency electrical stimulation (HFS, 100 pulses at 100 Hz, with each train interspaced by 20 s) to DIV 21 dissociated neuronal cultures (Fig. 5).

Ultrastructural analysis showed that NSBs become larger in size 5 minutes after stimulation (Fig. 5a,b): 8% increase in length (578 ± 14 nm from 534 ± 11 nm) and 17% increase in width (307 ± 7 nm from 263 ± 4 nm). The length of connectors was also slightly extended from 436 ± 14 nm without stimulation to 544 ± 19 nm while connector width did not change (125 ± 4 nm to 135 ± 3 nm, respectively). Fifteen minutes after stimulation NSB dimensions were similar to 5 min after stimulation (NSB length: 650 ± 15 nm, NSB width: 300 ± 6 nm) while connector dimensions were similar to unstimulated axons (conn. length: 410 ± 17 nm, conn.width:115 ± 4 nm).. These changes are consistent with the previously reported structural plasticity of axons in organotypic hippocampal slice cultures^11^, which persists for a long period following this type of tetanic stimulation. However, in our culture, this structural change was transient: over the next 30 minutes, the dimensions returned to the basal state (NSB length: 506 ± 11 nm, NSB width: 262 ± 6 nm, conn. length: 471 ± 13 nm, conn. width: 121 ± 3 nm) (Fig. 5a,b). Interestingly, similar changes in axon morphology were observed in neurons treated with 5 mM MßCD for 30 min (Fig. 5c,d). However, the resulting NSBs were smaller compared to those from the control (Fig. 5a), suggesting that modulation of the cholesterol level in the plasma membrane may be needed for induction of structural plasticity. Indeed, when we imaged the cholesterol level following HFS using NeonGreen-ALOD4 biosensors in wild-type neurons, the plasma membrane cholesterol level was decreased by ∼45% immediately after stimulation, and this reduction persisted over 30 min (Fig. 5e). These data suggest that this structural plasticity of axons may arise from changes in the plasma membrane cholesterol level.

To assess whether the axonal structural plasticity induces functional plasticity, we performed AP simulations based on the dimensions obtained in Fig. 5a-d. Our simulations predict that as axon nano-pearling dimensions increase, AP conduction velocity would decrease from 0.159 m/s to 0.152 m/s at 5’ then to 0.147m/s over 30 min in control neurons and remain unchanged over 30 min in MßCD-treated neurons (0.133 m/s to 0.143 m/s to 0.133 m/s, respectively) (Fig. 5g). Consistent with the modeling prediction, electrophysiology recordings in acute slices showed that the AP conduction velocity decreases upon HFS in control samples (baseline: 0.210 ± 0.07 m/s, 5’: 0.161 ± 0.04 m/s, 60’: 0.130 ± 0.03 m/s) but did not change in MβCD (baseline: 0.117 ± 0.01 m/s, 5’: 0.105 ± 0.02 m/s, 60’: 0.108 ± 0.01 m/s) (Fig. 5f,h). This change was long-lasting, persisting over 60 min, likely due to the expression of long-term potentiation in acute slices. Interestingly, when cholesterol is depleted from acute slices using MßCD, the conduction velocity did not change after high-frequency stimulation (Fig. 5f,h), suggesting that the cholesterol mobilization may be indeed needed for plasticity induction.

Together, these results suggest that neuronal stimulation modulates axon nano-pearling and function, in part, through the control of biophysical factors.

## Discussion

Axon morphology has been extensively studied over the last 70 years, revealing more and more complexity to axon structure. Our study uncovers further morphological complexity in that axons in the mammalian central nervous system under near-physiological conditions have a pearls-on-a-string morphology. Axon pearling is a well-characterized phenomenon that occurs even at the macroscopic level in neurons under stress^23, 54–59^. However, the morphology we describe here is on a nanoscale with the axon tract being only ∼60 nm in diameter with repeated varicosities with a diameter of 200 nm – the difference between the two regions is much below the diffraction limit of light. Thus, ultrastructural characterization is essential. However, the use of chemical fixatives for ultrastructural analysis leads to many artifacts in cell and tissue structure^60^ In fact, several studies using cryo-preservation techniques have noted a similar morphology in both myelinated and unmyelinated axons^22, 61^. In particular, Ochs *et al.* show that cat motor neurons exhibit pearled morphology *in vivo* when the limb of animals is straightened causing stretching of the nerve fibers and frozen by immersing the limb in liquid Freon-12.

Furthermore, intact Ctenophore^62^ and *Caenorhabditis elegans*^63^ neurons both exhibit axon nano-pearling, indicating that this nano-pearling is highly conserved. Thus, we propose that nano-pearled morphology is a ubiquitous and prominent feature of axons.

Why do axons pearl? Membrane mechanics studies describe how membrane pearling is caused by the energy minimization of homogeneous membrane tension along cylindrical membrane tubes. While axons do have complex structure, they are fundamentally membrane tubes and many of the same biophysical principles are applicable. Consistent with this notion, we found that NSB size is altered by the manipulation of in-plane membrane tension: increasing membrane fluidity or external osmotic pressure causes NSBs shrinkage. However, the connector regions are less affected by these treatments, suggesting that tension alone may not fully explain the axon morphology. Our results here indicate another component of membrane tension, the cytoskeleton. Given that perturbation of NMII and MTs results in the enlargement of the connector regions, the cytoskeleton appears to play an important role in determining connector morphology. Since NMII acts on F-actin, periodic actin rings may also contribute to morphology. However, axon morphology is unaltered when the MPS is disrupted with Latrunculin A or ßII spectrin KD, indicating that the MPS is not essential for axon pearling. Considering LatA treatment affects only a certain pool of F-actin^50, 64^, in the absence of actin rings NMII may still act on the remaining F-actin at the cortex under the LatA condition to maintain morphology. Nevertheless, it is important to note that the anchoring of the cytoskeleton to the plasma membrane is not as tight in the axon as in other cell types^65, 66^. Thus, cytoskeletal disruption may have a stronger impact in the connector region where the cytoskeleton is in closer contact with the plasma membrane. Finally, increasing membrane fluidity by depleting membrane cholesterol caused the NSBs to become rounder, showing that membrane properties also control axon nano-pearling. Together, these data suggest that axon nano-pearling is likely the result of membrane mechanics, particularly in-plane tension, with support from the cytoskeleton.

Axon nano-pearling has a strong implication for action potential propagation in unmyelinated axons. Previous work with cable theory modeling predicts that sudden changes in axon diameter, i.e. going from a thin connector to a large NSB, would slow AP propagation and at a certain size cause the AP propagation to fail^4, 5, 67, 68^. In agreement with this theory, our results here also suggest that the conduction velocity of APs is strongly dependent on axon geometry. In particular, the diameter of connectors has a linear relationship with velocity, as the cable theory predicts. However, the relationship between AP velocity and axon morphology is far more complicated. Seemingly, the ratio between the four parameters we measured is just as important as the absolute value. Intriguingly, there is an optimal NSB width/length ratio, ∼1.7, where action potential velocity is at its peak. Higher and lower than this value would result in slower AP conduction velocity. It is worth noting that the NSBs of neurons from acutely extracted brain tissue is at this ratio, while that of the NSBs of cultured neurons is about 2.

Thus, the surrounding physical environment can influence the morphology of axons and consequently, their function. Since mechanical properties are likely specific to each neuron type^69^, functional differences across various types of neurons may be attributed, at least in part, to differences in nano-pearled axon morphology. Further investigations are warranted.

One very intriguing idea that comes from our study of pearled morphology is that direct modulation of the biophysical forces and thereby axon morphology could, in turn, tune AP propagation velocity. In myelinated neurons, myelination placement and length are tightly regulated to control AP propagation, creating specific firing patterns important for various circuit functions such as coincident detection^70–72^. Through our modeling we can probe the effect of changing axon nano-pearling on AP propagation in unmyelinated axons. In treatments such as varying the osmolarity which cause a dramatic change in nano-pearling, there is a shift in AP propagation velocity. In fact, changing osmolarity has been shown to affect the burst-firing patterns of unmyelinated hippocampal CA1 neurons^73^, which may be due to the changes in axon nano-pearling. Likewise, Costas et al. (2020) has observed an increase in AP velocity from 0.4 m/s to 0.45m/s during NMII inhibition, which causes the actin ring diameter to increase from 300 nm to 400 nm. Finally, changes in membrane lipids have also been linked to changes in AP propagation. Recently, Korinek et al. (2020) found that cholesterol removal by MßCD disrupts the action potential propagation by causing action potential failure. Similar results are also obtained in crayfish neurons^74^. Likewise, our results here suggest that cholesterol depletion by MßCD slows down the AP propagation velocity. This effect may be due to the direct modulation of axon nano-pearling. However, cholesterol also plays an important role in channel clustering, and thus, the morphology may not be the sole contributor. Nonetheless, our work suggests a novel neuronal plasticity paradigm whereby modulation of biophysical factors controls axon nano-pearling and thereby, action potential conduction velocity.

## Methods

### Animals

The housing and care of all animals followed the NIH animal use guidelines and were approved by the Animal care and Use Committee at Johns Hopkins University School of Medicine.

C57/BL6J mice (wild-type) were used.

### Neuron culture

Both males and females were indistinguishably used in this study.

#### Astrocytes

Astrocytes were harvested from embryonic days 18 (E18)-postnatal day 0 (P0) cortices with trypsin treatment for 20 min at 37 °C with shaking, followed by dissociation and seeding on T-75 flask. Astrocytes were grown in full DMEM (DMEM (Gibco 10569010), 10% FBS (Thermo Fisher 26140079) and 100 U/mL penicillin-streptomycin (Gibco 15140122)) at 37 °C and 5% CO2 for 7 days. Two clean 6 mm sapphire disks (Technotrade Inc 616-100) were placed per well of 12- well tissue culture plate (Fisher Scientific 720081) and coated with poly-D-lysine (1 mg/ml, Sigma P6407) and collagen (Thermo Fisher Scientific A1048301). Astrocytes serve as a feeder layer for neurons and were seeded at 50,000/well one week before hippocampal neuronal culture. FUdR solution (DMEM, 0.81 mM FUdR (Sigma F0503), 2.04 mM Uridine (Sigma U3003)) was added to each well and incubated for 2-24 hrs at 37 °C before seeding neurons.

#### Neurons

E18 embryos were decapitated and stored in HBSS (Gibco 14175095). The hippocampi from each brain were removed in dissection media (1x HBSS, 100 U/mL penicillin-streptomycin, 1 mM pyruvate (Gibco 11360070), 10 mM Hepes (Gibco 15630080), 30 mM Glucose (Sigma G6152-100G)). Then, two hippocampi were placed in the same 15 ml tube with dissection media and 0.01% DNaseI (Sigma DN25) and 10 U/mL papain (Worthington LS003119) were added and incubated at 37 °C with gentle perturbation every 5 min for 20 min. The tissue was dissociated by gentle titration and run through a 70 μm cell strainer (Fisher scientific 22363548). Cells were spun down at 120x g for 2 min and gently resuspended in NM5 (Neurobasal medium (Gibco 21103049), 5% horse serum (Gibco 26050088), 100 U/mL penicillin/streptomycin, 2% GlutaMAX supplement (Gibco 35050061), 2% B27 (Gibco 17504044)). Cells were seeded on sapphire discs in 12-well plates at 75k cells/well in NM5.

The next day, the media was changed to NM0 culture media (Neurobasal medium, 2% GlutaMAX supplement, 1% B27). Half-media change was done at 14 days *in vitro* (DIV 14) with fresh NM0.

#### Organotypic hippocampal slice culture and freezing

Organotypic hippocampal slice cultures were performed using a protocol modified from Qian *et al* (2015). C57BL6 mouse pups were euthanized at P5-8 by rapid decapitation. Brains were harvested and dissected in ice-cold dissection medium (minimum essential medium, 24 mM HEPES, and 10 mM Tris-Cl). Forebrains were isolated and cut into 300–400 μm-thick coronal slices using a McIlwain tissue chopper (Ted Pella). Hippocampal slices were gently detached from the forebrain slices using forceps and placed onto Millicell inserts (30 mm diameter, 0.4 μm pore size). 3–4 slices were placed on a single insert. After carefully removing excessive liquid around the tissue slices, the inserts were placed into 6-well plates containing slice culture medium (50% MEM, 25% heat-inactivated horse serum, and 25% HBSS), with the membrane of the inserts just touching the surface of the medium. The slices were maintained in a 37 °C humidified incubator with 5% CO2. Medium in the plate was replaced with a low-serum medium (5% horse serum) at DIV1 and changed every two days from that point on.

#### STED in organotypic slice cultures

Organotypic hippocampal slices (Gähwiler type) from P5–7 wild-type mice were dissected and cultured 3–5 weeks in a roller drum at 35 °C^75^. Experimental procedures were in accordance with the European Union and CNRS UMR 5297 institutional guidelines for the care and use of laboratory animals (Council directive 2010/63/EU) and approved by the Committee of Ethics of Bordeaux (no. 50120198-A).

#### Acute cortical tissue extraction

Tissue for the neocortex of adult mice was prepared in the lab of Dr. Graham Knott with the ethics approved by the Swiss Federal Veterinary Office (the experimentation license 1889.3). Adult mice (C57 BL/6, 8 weeks old) were decapitated, and using scissors and forceps, the brain was immediately exposed. A piece of cortex was removed using the forceps and placed on the top of a closed plastic Petri dish containing ice. The tissue was then sliced with razor blades to produce 200 µm thick slices.

#### Chemical fixation

Adult mice (C57BL/6, 8 weeks old) were deeply anesthetized with inhalation anesthetic (isoflurane) and immediately perfused with a buffered solution of 2.5% glutaraldehyde, and 2% paraformaldehyde in phosphate buffer (0.1 M, pH 7.4, 250–300 ml per animal). One hour after perfusion, the brain was removed and 80-µm thick slices cut using a vibratome. These were washed in cacodylate buffer (0.1 M, pH 7.4, 3 × 5 min), post fixed in 1% osmium tetroxide and 1.5% potassium ferrocyanide in cacodylate buffer (0.1 M, pH 7.4, 40 min). They were then stained with 1% osmium tetroxide in cacodylate buffer (0.1 M, pH 7.4) for 40 min, and then in 1% uranyl acetate for 40 min before being dehydrated in a graded alcohol series, 3 min each change, and embedded in Durcupan resin. Specimens were cured at 60 °C for 24 hr.

### Drug treatments

#### Latrunculin A

Latrunculin A (Tocris Bioscience, 3973100U) was dissolved in DMSO to a stock concentration of 10 mM and then added to the cells for a final concentration of 20 µM and incubated in a cell incubator for the indicated time before the experiment. The stocks were used within one week.

#### Blebbistatin

(±)-Blebbistatin (Abcam, ab120425) was dissolved in DMSO to a stock concentration of 10 mM and then added to the cells for a final concentration of 10 µM and incubated for 1 hr in a cell incubator before the experiment. The stocks were used within one week.

#### Nocodazole

Nocodazole (Tocris Bioscience, 122810) was dissolved in DMSO to a stock concentration of 50 mM and then added to the cells for a final concentration of 50 µM and incubated for 1 hr in a cell incubator before the experiment. The stocks were used within one week.

#### Methyl-β-cyclodextrin

Methyl-β-cyclodextrin (MβCD) was dissolved directly in media for a final concentration of 5 mM and incubated for 30 min in a cell incubator before the experiment.

### Lentivirus production and infection

HEK-293T cells (1.2x10^6^ cells/flask, ATCC CRL-3216) were plated in T-75 flasks (Sarstedt 100437) coated with PDL (10 µg/ml) in full DMEM. T-75 flasks were incubated in 37 °C for 3 days. Cells were then resuspended with 0.05% Trypsin-EDTA (SigmaT1426) and replated at 6.5x10^6^ cells in 10 ml DMEM in a new PDL-coated T-75 flask. When cells were ∼90% confluent, the media was switched to NBA media (1% glutamax, 2% B27, and 0.2% penicillin-streptomycin). The modified shuttle vector (FUGW)^76^. containing expression constructs and helper plasmids (VSV-G and CMV-dR8.9) was mixed at 20, 5, and 7.5 µg, respectively, in 640 µl NaCl solution (150 mM) (solution I). Another solution (solution II) was prepared as follows: 246.7 µl H2O, 320 µl NaCl (300 mM), 73.3 µl polyethylenimine (0.6 µg/µl, Polysciences 24765- 2). Solutions I and II were combined and incubated at 24 °C for 10 min, followed by addition to the T-75 flask containing HEK293T cells. The cells were incubated at 37 °C (5% CO2) for 3 days. The media containing lentivirus was collected and virus particles were concentrated 20- fold using Amicon Ultracel-100k (EMD Millipore 901024). For all the experiments, dissociated hippocampal neurons were infected on DIV 9 with lentiviruses carrying the expression constructs.

### shRNA constructs

To express our shRNA in neurons, lentiviral expression constructs were used. All vectors were based on the lentiviral shuttle vector FUGW^76^. The ßII spectrin two sense sequences of shRNA are: 5′-GCATGTCACGATGTTACAA-3′ and 5′-GGATGAAATGAAGGTGCTA-3′ as previously described^50^ and were inserted using the Infusion HD Cloning kit (Takara Bio USA Inc, NC1470242).

### Viral infection for STED

For specifically labelling CA3 neurons, a glass micropipette backfilled with Sindbis-GFP viral particles diluted in TNE buffer containing (in M) 0.1 NaCl, 0.05 Tris-Cl (pH 8), 0.5 EDTA, and 0.001% Tween-20, connected to a pressure-injection device (Picospritzer, Parker, Hollis, NH, USA) was used. The pipette was positioned into CA3 pyramidal layer and the virus was injected by brief pressure pulses (50-150 ms; 10-15 psi) yielding an infection of approximately 30-50 neurons. Experiments were conducted 36-48 hrs after the infection giving sufficient GFP expression, while preserving the physiological health of the cells.

### Western Blot

Western blotting was used to verify knock-down efficacy. Protein lysates were obtained from cultures of hippocampal neurons infected with the shRNA containing lentiviral construct. Briefly, cells were lysed using RIPA lysis buffer (Pierce 89900). Proteins were separated by SDS-PAGE (BioRad 4561095) and transferred to nitrocellulose membrane (BioRad 1620115). Membranes were blocked with 5% milk and then incubated with either rabbit anti-β-actin (1:5000, SYSY 251003) or rabbit anti-GAPDH antibody (1:1000, Abcam ab37168) as a loading control and mouse anti-ßII spectrin (1:1000, Bd Cell Analysis BDB612563) antibodies overnight at 4 °C. Secondary antibody was added (Li-COR IRDye®, 800 cw, Goat anti-mouse 925-32210, 1:30,000 and IRDye® 680RD Goat anti-Rabbit IgG (H + L) 925-68071, 1:30,000 1 hr at room temperature) and results were imaged using a Li-COR ODYSSEY CLx (0958) and analyzed using the Image Studio Ver 5.2 software.

### Cholesterol sensor cloning and purification

Anthrolysin O domain 4 (ALOD4) tagged with NeonGreen was used to stain plasma membrane cholesterol of cultured hippocampal neurons in this study. ALOD4 (Addgene 111026) was N-terminally tagged with NeonGreen. Codon optimized NeonGreen was synthesized as gblock DNA from IDT. NeonGreen-ALOD4-His6 was cloned in pET28a vector backbone for bacterial expression. The plasmid was transformed into *E. coli* BL21 DE3 Rosetta cells, and protein expression was induced with 0.5 mM IPTG for 16 hrs at 4 °C. Histidine tagged NeonGreen- ALOD4 was purified using an AKTA purifier (GE). Briefly, cells were lysed using B-PER (Thermo 78243) cell lysis buffer supplemented with benzonase (25 U/ml), MgCl2 (2 mM) ATP (2 mM), imidazole (20 mM) and protease inhibitor (cOmplete, EDTA free, Roche). Cell debris was removed by centrifuging at 112,000 g for 20 min at 4 °C. The clear lysate was loaded onto 1 ml HisTrap FF (Cytiva) column pre-equilibrated with buffer A (20 mM HEPES (pH7.4), 150 mM NaCl, 20 mM imidazole). After washing the column with 10 volumes of buffer A, protein was eluted with a gradient of 20-500 mM imidazole. The purity of the protein was confirmed by SDS-PAGE, stained with Coomassie blue. More than 90% purity was achieved, and protein was stored in 20% glycerol as 40 µM stock solution.

### Cholesterol staining of neurons

To visualize cholesterol, hippocampal neurons (DIV21) were stained with NeonGreen-ALOD4 (1 µM final concentration) for 30 min at 37 °C. For depleting cholesterol from neuronal plasma membrane cells were treated with 5 mM MßCD (freshly dissolved as 6 mg powder per ml of NM0 media) and incubated for 30 min at 37 °C. MßCD containing media was removed by a quick wash with fresh media, 1 µM NeonGreen-ALOD4 was added to the cells and allowed to bind for 30 min at 37 °C. After the binding reaction, cells were briefly washed with media and fixed with 4% sucrose and 4% PFA in PBS and used for further antibody labeling.

For high frequency stimulation experiments and cholesterol labeling, neurons were cultured on grid coverslips (iBiDi, Grid 50). On DIV21, coverslips were placed in RC-BRFS slotted perfusion chamber with field stimulation (Warner Instruments), and cells were perfused with physiological saline solution (140 mM NaCl, 2.4 mM KCl, 10 mM HEPES, 10 mM Glucose (pH adjusted to 7.3 with NaOH), 4 mM CaCl2, and 1 mM MgCl2, 300 mOsm). The cells were stimulated with 3 trains of 100 pulses at 100 Hz with each train interspaced by 20 s, and either recovered for 30 min in a CO2 incubator at 37 °C or directly used for cholesterol labeling. For no stimulation control (sham), cells were placed in the perfusion chamber for the same time and recovered in media for 30 minutes. All treated cells were labeled with 1 µM NeonGreen-ALOD4 for 30 min.

### Immunofluorescence and Airyscan imaging

For Airyscan imaging, samples were imaged in Zeiss LSM880 (Carl Zeiss) in Airyscan mode. Fluorescence was acquired using a 63x objective lens (NA = 0.55) at 2048x2048 pixel size with the following settings: pixel dwell 1.02 ms and pin hole size above the lower limit for Airyscan imaging, as computed by ZEN software. For experiments comparing fluorescence intensities between no treatment and MßCD treated cells, staining and microscope settings were remained constant. NeonGreen-ALOD4 was used to stain cholesterol, and NeonGreen signal intensity was used to determine cholesterol level. Presynaptic regions were determined with either Synaptophysin or Bassoon. Axons were distinguished from dendritic processes based on their morphology, thin and no spines. 0.18 µm Z-sections were taken for each presynaptic bouton along axons. NeonGreen intensities, depicting plasma membrane cholesterol level in boutons, were quantitated in ImageJ.

For measuring cholesterol level in cells treated with or without MßCD, synaptophysin (SYSY, 101011, used at 1:100) was used to determine the synapses, and NeonGreen intensity was measured in ImageJ. The signal was normalized to the area of the selected axons. For high frequency stimulation experiments, axonal marker Ankyrin G (SYSY, 386004, used at 1:100) signal was used to determine the regions-of-interest (ROIs). All primary antibody incubations were performed overnight at 4 °C. All secondary antibodies were used at 1:500 for 45 min. For high frequency stimulation experiments, fluorescence intensity of cholesterol sensor was normalized to Ankyrin G signal and expressed as percentage of no treatment control.

### STED Microscopy

Note that the STED data used here are collected in the previous study^11^ and reanalyzed considering our findings on axon nano-pearling. For the detailed imaging method, please refer to Chéreau et al, 2017. Briefly, we used a home-built STED microscope based on an inverted microscope (DMI 6000 CS Trino; Leica), using a galvanometric beam scanner (Yanus IV; TILL Photonics) and a high numerical aperture objective lens (PL APO, 100X, oil, NA 1.4; Leica). The software Imspector (A. Schönle, MPI for Biophysical Chemistry) controlled image acquisition and parameters that minimize photodamage and photobleaching were chosen. All images were acquired with time-averaged powers of <6 μW for excitation and <8 mW for STED (measured at the back aperture of the objective). Image stacks were acquired with a voxel size of 19.5 nm (x, y), 375 nm (z) and a dwell time of 15 μs. A piezo z-focusing device (Physikinstrumente) controlled imaging depth, which was maximally 15 μm below tissue surface. Z-stacks of 40 × 40 × 3 μm (x, y, and z) were acquired every 6 min.

### High-pressure freezing and freeze-substitution

#### Dissociated cell culture

Cells cultured on sapphire disks were frozen using a high-pressure freezer (EM ICE, Leica Microsystems). Each disk with neurons was transferred into the physiological saline solution (140 mM NaCl, 2.4 mM KCl, 10 mM HEPES, 10 mM Glucose (pH adjusted to 7.3 with NaOH), 4 mM CaCl2, and 1 mM MgCl2, 300 mOsm) except in the cases of the hyper-osmotic and hypo-osmotic conditions which were 280 mM NaCl, 4.8 mM KCl, 20 mM HEPES, 20 mM Glucose, 4 mM CaCl2, and 1 mM MgCl2, 600 mOsm and 70 mM NaCl,1.2 mM KCl, 5 mM HEPES, 5 mM Glucose, 4 mM CaCl2, and 1 mM MgCl2, 150 mOsm respectively. Assembly was done in the freezing chamber, kept at 37 °C. The polycarbonate sample cartridges (Leica, 16771881, 16771882, & 16771838) were also warmed to 37 °C. Immediately after the sapphire disk was mounted on the sample holder and the cartridge was inserted into the freezing chamber. The disk was mounted onto the middle plate with neurons facing down into a specimen carrier, 100 µm side up (Technotrade, 610-100). A 200 µm spacer ring (Technotrade, 1259-100) was placed on top of the sapphire disk. The entire assembled middle plate was then placed on a piece of filter paper to remove the excess liquid, loaded between two half cylinders, and transferred into the freezing chamber. The frozen sample was automatically dropped into a storage Dewar filled with liquid nitrogen. After freezing, the middle plate with sapphire disks was transferred to a cup containing anhydrous acetone (−90 °C), which was placed in an automated freeze-substitution system (EM AFS2, Leica microsystems) using prechilled tweezers. Samples were transferred to specimen holders (Leica 1670715, 16707154) containing fixative (1% glutaraldehyde (EMS 16530), 0.1% Tannic acid (Sigma 403040-100G) in acetone), pre-cooled to -90 °C in the AFS. The freeze-substitution program was as follows: −90 °C for 44 hr, washed 5 times in pre-chilled acetone (30 min per wash), switched to 2% OsO4 (EMS 19132) in acetone, −90 °C for an additional 41 hr, −90 to −20 °C in 14 hr, −20 °C for 12 hr, and −20 to 4 °C in 2 hr.

### Organotypic slice culture

For high-pressure freezing experiments, the areas on the Millicell insert membrane where the hippocampal slices were placed were cut out with a scalpel. The slice along with the membrane at the bottom was soaked in a cryoprotectant (140 mM NaCl, 2.4 mM KCl, 10 mM HEPES, 10 mM glucose, 2 mM CaCl2, 3 mM MgCl2, and 20% bovine serum albumin) and placed onto a 6 mm sapphire disk. High-pressure freezing was performed as above using a Leica EM ICE device except freeze-substitution was performed in a fixative solution containing 1% OsO4, 1% methanol, 0.1% uranyl acetate, and 3% H2O dissolved in acetone. Samples were held at -90 °C for 24 hr, raised to -25 °C at a rate of 5 °C/hr, held at -25 °C for 12 hr, then raised to 0 °C at a rate of 5 °C/hr, held at 0 °C for 1 hr, and finally raised to 20 °C at a rate of 20 °C/hr. Samples were then washed with acetone, stained *en bloc* with 1% uranyl acetate, and washed again with acetone.

### Acutely extracted brain bits

Acutely extracted brain bits were placed inside 6 mm diameter aluminum sample holders with a 200-µm deep cavity. A drop of 1-hexadecene was added to ensure that no air was trapped once the sample holder was closed. This was high-pressure frozen using a Leica EM HPM100 (Leica Microsystems). The procedure was completed in less than 90 s from the moment of decapitation. The frozen samples were then stored in liquid nitrogen until further processing.

Frozen samples were embedded at low temperature (AFS2; Leica Microsystems). They were first exposed to 0.1% tannic acid in acetone, for 24 hr at −90°C, followed by 12 hr in 2% osmium tetroxide in acetone at the same temperature. The temperature was then raised to −30 °C over 72 hr and then the liquid replaced with pure acetone and the temperature increased to −10 °C over 24 hr. Finally, the tissue samples were mixed with increasing concentrations of epon resin over 8 hr whilst the temperature rose to 20 °C. They were then added to 100% resin for 2 hr and then placed in silicon molds for 24 hr at 65 °C for the resin to harden.

For both high-pressure frozen and chemically fixed acute tissue in resin, serial sections (50–100 sections per ribbon) from the tissue samples were collected on single slot copper grids with a pioloform support film. Images were collected in series using a transmission electron microscope (Tecnai Spirit, FEI Company) operating at 80 kV, housing a digital camera (FEI Eagle 4k × 4k).

### Embedding and sectioning

Following freeze-substitution, samples were washed with anhydrous acetone 4 times, 10 min each at room temperature. After washing, samples were infiltrated through 30%, and 70%, epon araldite (epon 6.2 g, araldite 4.4 g, DDSA 12.2 g, and BDMA 0.8 ml (Ted Pella 18012) in anhydrous acetone every 2 hr. Then, samples were transferred to caps of polyethylene BEEM capsules (EMS 102096-558) with 90% epon araldite and incubated overnight at 4 °C. Next day, samples were moved into new caps with fresh 100% epon araldite every 1 hr, 3 times, after which resins were cured at 60 °C for 48 hr. Once cured, sapphire disks were removed from resin, leaving the cells behind. Then, the block was cut and fixed onto a dummy block using super glue for sectioning. 40-nm sections were cut using an ultramicrotome (EM UC7, Leica microsystems) and collected on single-slot copper grids (Ted Pella 1GC12H) coated with 0.7% pioloform in chloroform for transmission electron microscopy imaging or on silicon wafers. The sections were stained with 2.5% uranyl acetate (Ted Pella, 19481) in 75% methanol.

### Transmission electron microscopy

Samples were imaged at 80 kV at the 20,000x magnification on a Hitachi 7600 transmission electron microscope equipped with a dual AMT CCD camera system. Images were acquired through AMT Capture v6. Each sample was given a random number and about 50 electron micrographs per sample were taken.

### Array tomography

The serial sections were imaged using a scanning electron microscope (SEM), a JSM-7900F (JEOL Ltd., Tokyo, Japan), with a backscattered electron (BSE) detector. These sections were imaged automatically with Array Tomography Supporter, a custom software for serial-section SEM, also called array tomography^77^, at an acceleration voltage of 7 kV with a resolution of 2.0 nm per pixel.

### Electron microscopy image analysis

Each set of images from a single experiment were shuffled for analysis as a single pool using a custom R (R Development Team, R Studio 1.3, R version 3.5.1) script. Images that did not contain an axon or could not be reliably segmented were excluded from segmentation after images were randomized. No other data were excluded. The NSBs and connectors were measured in Fiji (version 1.0) using a custom macro. Any boutons containing 40 nm diameter vesicles were excluded as synaptic boutons. To minimize any possible bias and to maintain consistency, all image segmentation, still in the form of randomized files, were checked by a second lab member. Dimensions were then quantitated using custom MATLAB (MathWorks R2017-R2020a) scripts (available from: https://github.com/ shigekiwatanabe/SynapsEM). For 3D data, dimensions were determined as follows: first, dimensions in 2D were calculated as above, then 3D dimensions were calculated using the Pythagorean theorem with the assumption that each image was aligned and each section is of equal thickness. All example micrographs shown had their brightness and contrast adjusted to be similar depending on the raw image, rotated, and cropped in Adobe Illustrator.

### Electrophysiology

Adult mice of both sexes ranging from 6-8 weeks of age were anaesthetized using a combination of Isoflurane inhalation and Avertin injection. Mice underwent cardiac perfusion using chilled sucrose solution (10 mM NaCl, 2.5 mM KCl,10 mM Glucose, 84 mM NaHCO3, 120 mM NaH2PO4, 195 mM Sucrose, 1 mM CaCl2, 2 mM MgCl2) saturated with oxygen 5% / carbon dioxide 95% (carbogen). Brain was rapidly dissected, and hippocampi removed. Hippocampi were then embedded in premade agarose molds and sliced at 400 µm using Leica VT1200S vibratome at a speed of 0.05 mm/s and amplitude of 1.0 mm. Slices were then transferred to artificial cerebrospinal fluid (ACSF; 119 mM NaCl, 2.5 mM KCl, 1.3 mM MgSO4, 2.5 mM CaCl2, 26 mM NaHCO3, 1 mM NaH2PO4, and 11 mM D-glucose (315 Osm, pH 7.4) heated using a water bath to 32 °C saturated with carbogen. Slices were recovered at this temperature for 15 min before being removed from the bath and recovered for 1 hour at room temperature.

Recordings were performed at 32 °C in ACSF containing 2 mg/ml Kynurenic acid to suppress excitatory postsynaptic potential. Slices treated with MβCD (5 mM in ACSF) were pre-incubated 30 minutes prior to recording. MβCD was also present in recording ACSF. Glass pipettes containing silver chloride electrodes were used to both stimulate and record. The stimulating electrode was filled with ACSF and placed in CA1 to stimulate the Schaffer collaterals. Recording electrode was filled with 1M NaCl and placed at varying distances ranging from 0.2-1 mm away from the stimulating electrode towards CA2/3 in CA1 Schaffer collaterals. A bipolar square pulse of 0.3 ms at 60 mV was applied every 1 minute for 20 minutes and averaged to determine baseline. Long term potentiation was induced through applying three trains of 100 pulses at 100 Hz separated by a 20 s interval. A bipolar square pulse of 0.3 ms at 60 mV was applied every 1 min for 1 hr after LTP induction. Recordings were taken using Multiclamp 700B and Digidata 1550B and Clampex v11.2 software. Stimulus was applied using A-M Systems Isolated Pulse Stimulator Model 2100. 1-3 slices per condition per mouse were recorded. Traces were analyzed using Clampfit software v11.2 and fiber volley speed was determined (time to peak/distance traveled).

### Statistical analysis

All distributions shown of NSBs and connectors are pooled from multiple experiments. All data were initially examined on a per experiment basis (within a freezing done on the same day and all segmentation done in a single randomized batch). We did not predetermine sample sizes using power analysis, but based them (N= 2 and 3, n > 100) on our prior studies^78–82^. All data were tested for normality by the D’Agostino–Pearson omnibus test to determine whether to use parametric or nonparametric methods. Comparisons between multiple groups followed by full pairwise comparisons were performed using one-way analysis of variance followed by Kruskal–Wallis test followed by Dunn’s multiple comparisons test. All statistical analyses were performed and all graphs created in Graphpad Prism.

### Morphology simulations

The procedures are described in Supplementary Information.

### AP modeling

The procedures are described in Supplementary Information.

### Data availability

All original data are available through Figshare.com or upon request. All original codes are archived on Zenodo and available from Github or upon request.

## Author Contributions

J.G., M.B.Q., R.P., C.L., P.R., and S.W. conceived the study and designed the experiments. P.R. and S.W. oversaw the overall projects and funded the research. J.G., S.M., S.S., performed ultrastructural analysis. M.B.Q., C.L., C.Z., M.B. performed modeling. Q.G., Y.Y., and M.S. provided array tomography data from organotypic slice cultures. R.C. and V.N. provided STED data. G.K. provided images from acutely extracted mouse brain tissues. S.R. performed fluorescence imaging of cholesterol biosensors. Co-second authors M.B.Q., R.P., and C.L. contributed equally, and the order is simply alphabetical in reverse.

## Materials and Correspondence

All correspondence should be addressed to Padmini Rangamani and Shigeki Watanabe.

---

**Extended Data Fig. 1**. related to Fig. 1. Axon is pearled, not tubular, under homeostatic conditions.

a. Example micrographs showing acutely extracted mouse brain tissue either chemically fixed (left) or high-pressure frozen (right). The boxed axons are reconstructed using IMOD. Note that chemically fixed axons are like cylindrical tubes. Scale bar: 500 nm.

b. Higher magnification images of the boxed area in a. Scale bar: 200 nm.

c. Additional representative images of axons from acutely extracted brain tissue, organotypic slice culture, and dissociated neuron culture. Scale bar: 200 nm.

d. Super Plots of Fig. 1d, showing experimental variability. Median and 95% confidence intervals are shown. N = 3 and n = 30 axons from each acutely extracted sample, n = 133 axons from the organotypic sample, N = 3 and n = 100 axons from each dissociated sample. Each color represents one replicate. Each dot is one axon.

e. Additional STED micrographs showing pearled axon morphology in live neurons from organotypic slice cultures. Scale bar: 500 nm. Individual axons are straightened and represented on the right panels. Scale bar: 200 nm.

**Extended Data Fig. 2**, related to Fig. 2.

a. Axon morphology is modeled using classic Helfrich membrane model and governed by the membrane bending, surface tension, and osmotic condition. 3D model prediction of axon morphology in indicated osmotic conditions, tension of 0.001 mN/m, and bending rigidity of 50kBT. Scale bar: 200 nm.

b. 3D model prediction of axon morphology at indicated bending rigidity, tension of 0.001 mN/m, and osmolarity 300 mOsm. Scale bar: 200 nm.

c. Rotationally averaged shape profiles of axons under constant 0.001 mN/m tension and varying osmotic conditions and bending rigidity. The bead diameter (D) and length (L) in nanometers are given as a text inset for each condition.

d. Rotationally averaged shape profiles of axons under constant 0.01 mN/m tension and varying osmotic conditions and bending rigidity. The bead diameter (D) and length (L) in nanometers are given as a text inset for each condition.

**Extended Data Fig. 3**, related to Fig. 2

a. Additional example micrographs of axons high-pressure frozen at indicated osmotic conditions. Scale bar: 200 nm.

b. Super Plots of Fig. 2f, showing experimental variability. Median and 95% confidence intervals are shown. N = 3 independent cultures and n = 100 axons each. Each color represents one replicate. Each dot is one axon.

c. Example micrographs of axons high-pressure frozen at indicated osmotic conditions. Mannitol was used to adjust the osmolarity. Scale bar: 200 nm.

d. Super Plots of Extended Data Fig. 2e, showing the dimensions of NSBs at indicated osmotic conditions. Median and 95% confidence intervals are shown. N = 3 independent cultures and n = 100 axons each. Each color represents one replicate. Each dot is one axon.

e. Plot showing the dimensions of NSBs at indicated osmotic conditions. Mean and SEM are plotted. N = 3 independent cultures, n = 100 axons each.

*n.s*. not significant, *\*\** p<0.01, *\*\*\*\** p<0.0001.

**Extended Data Fig. 4**. related to Fig. 2

a. NeonGreen-ALOD4 staining of cultured mouse hippocampal neurons treated with sham (control) or 5 mM MßCD for 30 min. Scale bar = 5 µm.

b. Plot showing the normalized intensity of NeonGreen-ALOD4. Signals are normalized by the area of each region-of-interest. Control: 2773 ± 234 A.U., MβCD: 86 ± 13 A.U., median and 95% confidence intervals are plotted. A.U. arbitrary unit.

c. Additional example micrographs of axons high-pressure frozen at indicated osmotic conditions. Scale bar: 200 nm.

d. Super Plots of Fig. 2h, showing variability. N = 3 independent cultures and n = 100 axons each median and 95% confidence intervals are plotted. *n.s.* not significant, \*\*\**P* < 0.001, **** *P* < 0.0001. Each color represents one replicate. Each dot is one axon.

**Extended Data Fig. 5**, related to Fig. 3

a. Western blots and quantification showing the efficiency of shRNA-mediated knock down (KD) of ßII spectrin. Scr, scramble.

b. Additional example micrographs of axons from neurons treated with indicated drugs. Scale bar: 200 nm.

c. Super Plots of Fig. 3a, showing experimental variability. Median and 95% confidence intervals are shown. N = 3 independent cultures and n = 100 axons each. Each color represents one replicate. Each dot is one axon.

d. Additional example micrographs of axons from neurons treated with indicated drugs. Note that these neurons are treated with DMSO or Latrunculin A for 1 hour. Scale bar: 200 nm.

e. Super Plots, showing experimental variability of d. Median and 95% confidence intervals are shown. N = 1 and n = 100 axons. Each dot is one axon.

f. Additional example micrographs of axons from neurons infected with lentivirus carrying scramble shRNA or shRNA against ßII spectrin. Scale bar: 200 nm.

g. Super Plots of Fig. 3d, showing experimental variability. Median and 95% confidence intervals are shown. N = 3 independent cultures and n = 100 axons each. Each color represents one replicate. Each dot is one axon.

h. Additional example micrographs of axons from neurons treated with indicated drugs. Scale bar: 200 nm.

i. Super Plots of Fig. 3f, showing experimental variability. Median and 95% confidence intervals are shown. N = 3 independent cultures and n = 100 axons each.

*n.s.*, not significant, \*\**P* < 0.01,\*\*\**P* < 0.001, \*\*\*\**P* < 0.0001.

**Extended Data Fig. 6**, related to Fig. 4

a-f, Plots showing the relationship between AP velocity and the ratio of two measured values like connector width/NSB width (a), connector width/NSB length (b), connector width/connector length (c), connector length/NSB length (d), NSB length/NSB width (e), and connector length/NSB width (f). The data are fitted by a simple linear regression in all except for e, which is fitted by a gaussian non-linear curve.

**Extended Data Fig. 7**, related to Fig.5

a. Additional example micrographs showing axon morphology from neurons unstimulated or stimulated with 3x100 pulses at 100 Hz (high-frequency stimulation, HFS) and high-pressure frozen at 5 min and 30 min after stimulation. Scale bar: 200 nm

b. Super Plots of Fig. 5b, showing experimental variability. Median and 95% confidence intervals are shown. N = 3 independent cultures except for control 15’ post stim and MβCD 5’ post stim which were N = 2. n = 100 axons each. Each color represents one replicate. Each dot is one axon. *n.s.*, not significant, \**P* < 0.05, \*\**P* < 0.01,\*\*\**P* < 0.001, \*\*\*\**P* < 0.0001

## References

1. Hodgkin, A. L. & Huxley, A. F. A quantitative description of membrane current and its application to conduction and excitation in nerve. J. Physiol. 117, 500–544 (1952).

2. Waxman, S. G. Determinants of conduction velocity in myelinated nerve fibers. Muscle Nerve 3, 141–150 (1980).

3. Acker, C. D. & White, J. A. Roles of IA and morphology in action potential propagation in CA1 pyramidal cell dendrites. J. Comput. Neurosci. 23, 201–216 (2007).

4. Zhou, Y. & Bell, J. Study of propagation along nonuniform excitable fibers. Math. Biosci. 119, 169–203 (1994).

5. Goldstein, S. S. & Rall, W. Changes of action potential shape and velocity for changing core conductor geometry. Biophys. J. 14, 731–757 (1974).

6. Waxman, S. G. & Bennett, M. V. I. Relative Conduction Velocities of Small Myelinated and Non-myelinated Fibres in the Central Nervous System. Nat. New Biol. 1972 23885 **238**, 217–219 (1972).

7. Iwasa, K., Tasaki, I. & Gibbons, R. C. Swelling of Nerve Fibers Associated with Action Potentials. Science (80-.). 210, 338–339 (1980).

8. Hill, B. C., Schubert, E. D., Nokes, M. A. & Michelson, R. P. Laser interferometer measurement of changes in crayfish axon diameter concurrent with action potential. Science 196, 426–428 (1977).

9. Tasaki, I. & Byrne, P. M. Swelling of frog dorsal root ganglion and spinal cord produced by afferent volley of impulses. Brain Res. 272, 360–363 (1983).

10. Tasaki, I. & Byrne, P. M. Large mechanical changes in the bullfrog olfactory bulb evoked by afferent fiber stimulation. Brain Res. 475, 173–176 (1988).

11. Chéreau, R., Saraceno, G. E., Angibaud, J., Cattaert, D. & Nägerl, U. V. Superresolution imaging reveals activity-dependent plasticity of axon morphology linked to changes in action potential conduction velocity. Proc. Natl. Acad. Sci. U. S. A. 114, 1401–1406 (2017).

12. Shepherd, G. M. G. & Harris, K. M. Three-Dimensional Structure and Composition of CA3→CA1 Axons in Rat Hippocampal Slices: Implications for Presynaptic Connectivity and Compartmentalization. J. Neurosci. 18, 8300–8310 (1998).

13. Alimohamadi, H. & Rangamani, P. Modeling membrane curvature generation due to membrane–protein interactions. Biomolecules vol. 8 (2018).

14. Bar-Ziv, R., Tlusty, T. & Moses, E. Critical dynamics in the pearling instability of membranes. Phys. Rev. Lett. 79, 1158–1161 (1997).

15. Bar-Ziv, R. & Moses, E. Instability and ‘pearling’ states produced in tubular membranes by competition of curvature and tension. Phys. Rev. Lett. 73, 1392–1395 (1994).

16. Naito, H., Okuda, M. & Zhong-Can, O. Y. New Solutions to the Helfrich Variation Problem for the Shapes of Lipid Bilayer Vesicles: Beyond Delaunay’s Surfaces. Phys. Rev. Lett. 74, 4345 (1995).

17. Campelo, F. & Hernández-Machado, A. Model for curvature-driven pearling instability in membranes. Phys. Rev. Lett. 99, 088101 (2007).

18. Heinrich, D., Ecke, M., Jasnin, M., Engel, U. & Gerisch, G. Reversible Membrane Pearling in Live Cells upon Destruction of the Actin Cortex. Biophys. J. 106, 1079–1091 (2014).

19. Pullarkat, P. A., Dommersnes, P., Fernández, P., Joanny, J.-F. & Ott, A. Osmotically Driven Shape Transformations in Axons. Phys. Rev. Lett. 96, 048104 (2006).

20. Yuan, F. et al. Membrane bending by protein phase separation. Proc. Natl. Acad. Sci. U.S. A. 118, e2017435118 (2021).

21. Alimohamadi, H., Ovryn, B. & Rangamani, P. Modeling membrane nanotube morphology: the role of heterogeneity in composition and material properties. Sci. Rep. 10, 1–15 (2020).

22. Ochs, S. & Jersild, R. A. Cytoskeletal organelles and myelin structure of beaded nerve fibers. Neuroscience 22, 1041–1056 (1987).

23. Datar, A. et al. The Roles of Microtubules and Membrane Tension in Axonal Beading, Retraction, and Atrophy. Biophys. J. 117, (2019).

24. Moor, H. Theory and Practice of High Pressure Freezing. Cryotech. Biol. Electron Microsc. 175–191 (1987) doi:10.1007/978-3-642-72815-0_8.

25. Studer, D., Humbel, B. M. & Chiquet, M. Electron microscopy of high pressure frozen samples: bridging the gap between cellular ultrastructure and atomic resolution. Histochem. Cell Biol. 130, 877–889 (2008).

26. Hoffman, D. P. et al. Correlative three-dimensional super-resolution and block face electron microscopy of whole vitreously frozen cells. Science 367, (2020).

27. Westrum, L. E. & Blackstad, T. W. An electron microscopic study of the stratum radiatum of the rat hippocampus (regio superior, CA 1) with particular emphasis on synaptology. J. Comp. Neurol. 119, 281–309 (1962).

28. Major, G. et al. Detailed Passive Cable Models of Whole-Cell Recorded CA3 Pyramidal Neurons in Rat Hippocampal Slices. J. Neurosci. 4, 4613–4638 (1994).

29. Shepherd, G. M. G., Raastad, M. & Andersen, P. General and variable features of varicosity spacing along unmyelinated axons in the hippocampus and cerebellum. Proc. Natl. Acad. Sci. U. S. A. 99, 6340–6345 (2002).

30. Debanne, D. Information processing in the axon. Nat. Rev. Neurosci. 2004 54 **5**, 304–316 (2004).

31. Bucher, D. & Goaillard, J. M. Beyond faithful conduction: Short-term dynamics, neuromodulation, and long-term regulation of spike propagation in the axon. Prog. Neurobiol. 94, 307–346 (2011).

32. Tamada, H., Blanc, J., Korogod, N., Petersen, C. C. H. C. & Knott, G. W. Ultrastructural comparison of dendritic spine morphology preserved with cryo and chemical fixation. Elife 9, 1–15 (2020).

33. Grevesse, T., Dabiri, B. E., Parker, K. K. & Gabriele, S. Opposite rheological properties of neuronal microcompartments predict axonal vulnerability in brain injury. Sci. Reports 2015 51 **5**, 1–10 (2015).

34. Korogod, N., Petersen, C. C. H. & Knott, G. W. Ultrastructural analysis of adult mouse neocortex comparing aldehyde perfusion with cryo fixation. Elife 4, (2015).

35. Helfrich, W. Elastic Properties of Lipid Bilayers: Theory and Possible Experiments. Zeitschrift fur Naturforsch. - Sect. C J. Biosci. 28, 693–703 (1973).

36. Canham, P. B. The minimum energy of bending as a possible explanation of the biconcave shape of the human red blood cell. J. Theor. Biol. 26, 61–81 (1970).

37. Deserno, M., Kremer, K., Paulsen, H., Peter, C. & Schmid, F. Computational Studies of Biomembrane Systems: Theoretical Considerations, Simulation Models, and Applications. 237–283 (2013) doi:10.1007/12_2013_258.

38. The Role of Mechanics in the Study of Lipid Bilayers. 577, (2018).

39. Rangamani, P. The many faces of membrane tension: Challenges across systems and scales. Biochim. Biophys. Acta - Biomembr. 1864, 183897 (2022).

40. Zhu, C., Lee, C. T. & Rangamani, P. Mem3DG: Modeling membrane mechanochemical dynamics in 3D using discrete differential geometry. Biophys. Reports 2, (2022).

41. Tsafrir, I. et al. Pearling Instabilities of Membrane Tubes with Anchored Polymers. Phys. Rev. Lett. 86, 1138 (2001).

42. Su, Y. C. & Chen, J. Z. Y. A model of vesicle tubulation and pearling induced by adsorbing particles. Soft Matter 11, 4054–4060 (2015).

43. Wasser, C. R., Ertunc, M., Liu, X. & Kavalali, E. T. Cholesterol-dependent balance between evoked and spontaneous synaptic vesicle recycling. J. Physiol. 579, 413 (2007).

44. Korinek, M. et al. Cholesterol modulates presynaptic and postsynaptic properties of excitatory synaptic transmission. Sci. Rep. 10, 12651 (2020).

45. Gay, A., Rye, D. & Radhakrishnan, A. Switch-like Responses of Two Cholesterol Sensors Do Not Require Protein Oligomerization in Membranes. Biophys. J. 108, 1459–1469 (2015).

46. Diz-Muñoz, A., Fletcher, D. A. & Weiner, O. D. Use the force: membrane tension as an organizer of cell shape and motility. Trends Cell Biol. 23, 47–53 (2013).

47. Venkova, L. et al. A mechano-osmotic feedback couples cell volume to the rate of cell deformation. Elife 11, (2022).

48. De Belly, H. et al. Cell protrusions and contractions generate long-range membrane tension propagation. Cell 0, 1–13 (2023).

49. Xu, K., Zhong, G. & Zhuang, X. Actin, spectrin, and associated proteins form a periodic cytoskeletal structure in axons. Science 339, 452–6 (2013).

50. Zhong, G. et al. Developmental mechanism of the periodic membrane skeleton in axons. Elife 3, (2014).

51. Costa, A. R. C. et al. The membrane periodic skeleton is an actomyosin network that regulates axonal diameter and conduction. Elife 9, (2020).

52. Spector, I., Braet, F., Shochet, N. R. & Bubb, M. R. New anti-actin drugs in the study of the organization and function of the actin cytoskeleton. Microsc. Res. Tech. 47, 18–37 (1999).

53. Koch, C. Biophysics of Computation: Information Processing in Single Neurons. Biophys. Comput. (1998) doi:10.1093/OSO/9780195104912.001.0001.

54. Yagishita, S. Morphological investigations on axonal swellings and spheroids in various human diseases. Virchows Arch. A Pathol. Anat. Histol. 378, 181–197 (1978).

55. Büki, A., Okonkwo, D. O., Wang, K. K. W. & Povlishock, J. T. Cytochrome c Release and Caspase Activation in Traumatic Axonal Injury. J. Neurosci. 20, 2825–2834 (2000).

56. El-Khodor, B. F. & Burke, R. E. Medial forebrain bundle axotomy during development induces apoptosis in dopamine neurons of the substantia nigra and activation of caspases in their degenerating axons. J. Comp. Neurol. 452, 65–79 (2002).

57. Kerschensteiner, M., Schwab, M. E., Lichtman, J. W. & Misgeld, T. In vivo imaging of axonal degeneration and regeneration in the injured spinal cord. Nat. Med. 2005 115 **11**, 572–577 (2005).

58. Stokin, G. B. et al. Axonopathy and transport deficits early in the pathogenesis of Alzheimer’s diseases. Science *(80-.).* **307**, 1282–1288 (2005).

59. Tagliaferro, P. & Burke, R. E. Retrograde Axonal Degeneration in Parkinson Disease. J. Parkinsons. Dis. 6, 1–15 (2016).

60. Korogod, N., Petersen, C. C. H. & Knott, G. W. Ultrastructural analysis of adult mouse neocortex comparing aldehyde perfusion with cryo fixation. Elife 4, (2015).

61. Ma, D., Deng, B., Sun, C., McComb, D. W. & Gu, C. The Mechanical Microenvironment Regulates Axon Diameters Visualized by Cryo-Electron Tomography. Cells 11, (2022).

62. Burkhardt, P. et al. Syncytial nerve net in a ctenophore adds insights on the evolution of nervous systems. Science *(80-.).* **380**, 293–297 (2023).

63. Jurrus, E. et al. Semi-automated Neuron Boundary Detection and Nonbranching ProcessSegmentation in Electron Microscopy Images. Neuroinformatics 11, 5 (2013).

64. Abouelezz, A., Micinski, D., Lipponen, A. & Hotulainen, P. Sub-membranous actin rings in the axon initial segment are resistant to the action of latrunculin. Biol. Chem. 400, 1141– 1146 (2019).

65. Datar, A., Bornschlögl, T., Bassereau, P., Prost, J. & Pullarkat, P. A. Dynamics of membrane tethers reveal novel aspects of cytoskeleton-membrane interactions in axons. Biophys. J. 108, 489–497 (2015).

66. Shi, Z., Innes-Gold, S. & Cohen, A. E. Membrane tension propagation couples axon growth and collateral branching. Sci. Adv. 8, 31 (2022).

67. Altenberger, R., Lindsay, K. A., Ogden, J. M. & Rosenberg, J. R. The interaction between membrane kinetics and membrane geometry in the transmission of action potentials in non-uniform excitable fibres: a finite element approach. J. Neurosci. Methods 112, 101–117 (2001).

68. Maia, P. D. & Kutz, J. N. Identifying critical regions for spike propagation in axon segments. J. Comput. Neurosci. 36, 141–155 (2014).

69. Magdesian, M. H. et al. Atomic Force Microscopy Reveals Important Differences in Axonal Resistance to Injury. Biophys. J. 103, 405–414 (2012).

70. Salami, M., Itami, C., Tsumoto, T. & Kimura, F. Change of conduction velocity by regional myelination yields constant latency irrespective of distance between thalamus and cortex. Proc. Natl. Acad. Sci. U. S. A. 100, 6174–6179 (2003).

71. Lang, E. J. & Rosenbluth, J. Role of myelination in the development of a uniform olivocerebellar conduction time. J. Neurophysiol. 89, 2259–2270 (2003).

72. Seidl, A. H. & Rubel, E. W. Systematic and differential myelination of axon collaterals in the mammalian auditory brainstem. Glia 64, 487–494 (2015).

73. Azouz, R., Alroy, G. & Yaari, Y. Modulation of endogenous firing patterns by osmolarity in rat hippocampal neurones. J. Physiol. 502, 175–187 (1997).

74. Zamir, O. & Charlton, M. P. Cholesterol and synaptic transmitter release at crayfish neuromuscular junctions. J. Physiol. 571, 83–99 (2006).

75. Tønnesen, J., Katona, G., Rózsa, B. & Nägerl, U. V. Spine neck plasticity regulates compartmentalization of synapses. Nat. Neurosci. 2014 175 **17**, 678–685 (2014).

76. Lois, C., Hong, E. J., Pease, S., Brown, E. J. & Baltimore, D. Germline transmission and tissue-specific expression of transgenes delivered by lentiviral vectors. Science (80-.). 295, 868–872 (2002).

77. Micheva, K. D. & Smith, S. J. Array Tomography: A New Tool for Imaging the Molecular Architecture and Ultrastructure of Neural Circuits. Neuron 55, 25–36 (2007).

78. Watanabe, S. et al. Ultrafast endocytosis at mouse hippocampal synapses. Nature 504, 242–247 (2013).

79. Watanabe, S. et al. Ultrafast endocytosis at Caenorhabditis elegans neuromuscular junctions. Elife 2013, (2013).

80. Watanabe, S. et al. Clathrin regenerates synaptic vesicles from endosomes. Nature 515, 228–233 (2014).

81. Kusick, G. F. et al. Synaptic vesicles transiently dock to refill release sites. Nat. Neurosci. 23, 1329–1338 (2020).

82. Li, S. et al. Asynchronous release sites align with NMDA receptors in mouse hippocampal synapses. Nat. Commun. 2021 121 **12**, 1–13 (2021).

