## Extended Data Fig. for "Membrane mechanics dictate axonal morphology and function"

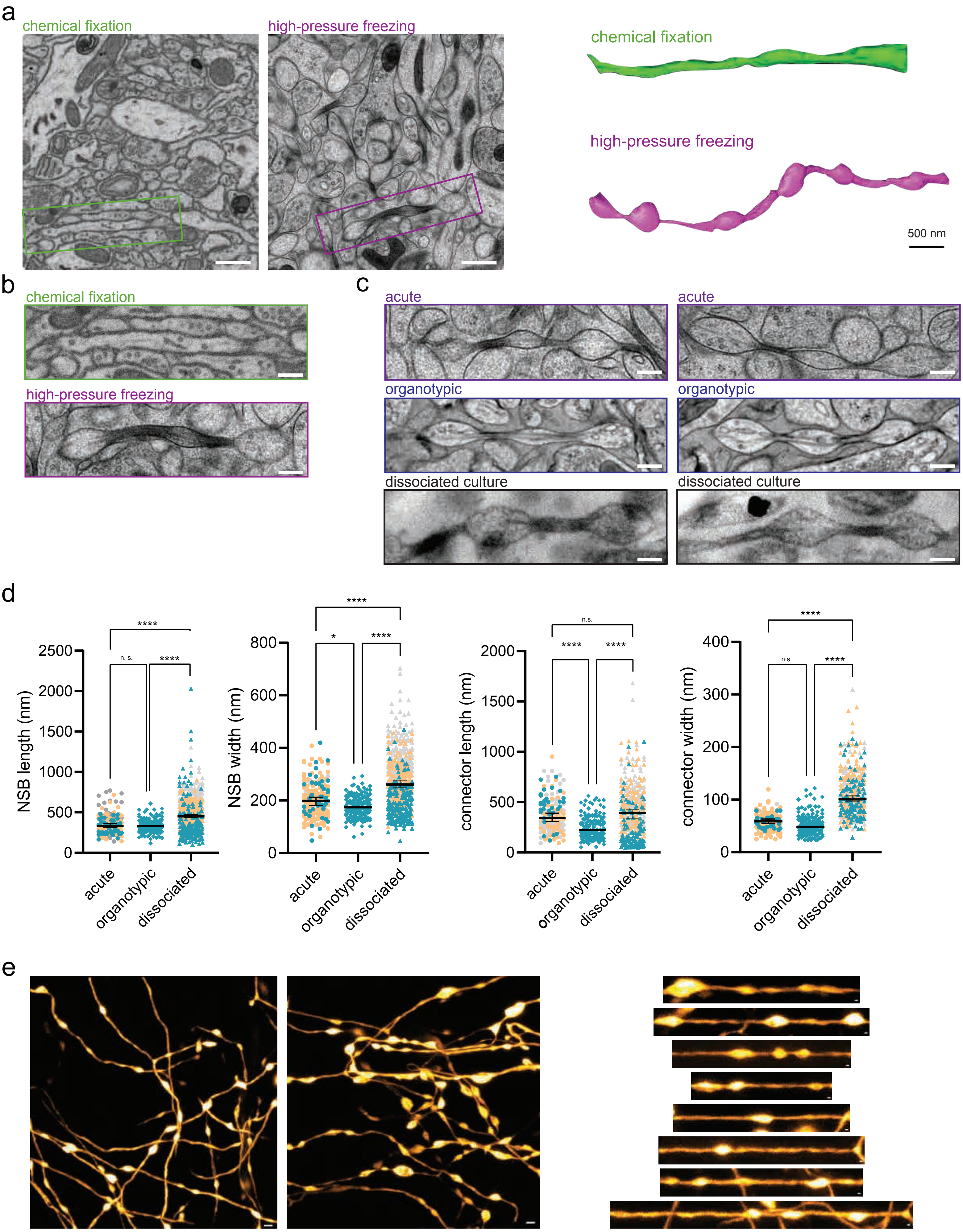
a
chemical fixation
high-pressure freezing
chemical fixation
high-pressure freezing
500 nm
b
chemical fixation
high-pressure freezing
c
acute
acute
organotypic
organotypic
dissociated culture
dissociated culture
d
NSB length (nm)
NSB width (nm)
connector length (nm)
connector width (nm)
acute
organotypic
dissociated
****
n. s.
*
n.s.
e

**Extended Data Fig. 1**. related to Fig. 1. Axon is pearled, not tubular, under homeostatic conditions.

a. Example micrographs showing acutely extracted mouse brain tissue either chemically fixed (left) or high-pressure frozen (right). The boxed axons are reconstructed using IMOD. Note that chemically fixed axons are like cylindrical tubes. Scale bar: 500 nm.

b. Higher magnification images of the boxed area in a. Scale bar: 200 nm.

c. Additional representative images of axons from acutely extracted brain tissue, organotypic slice culture, and dissociated neuron culture. Scale bar: 200 nm.

d. Super Plots of Fig. 1d, showing experimental variability. Median and 95% confidence intervals are shown. N = 3 and n = 30 axons from each acutely extracted sample, n = 133 axons from the organotypic sample, N = 3 and n = 100 axons from each dissociated sample. Each color represents one replicate. Each dot is one axon.

e. Additional STED micrographs showing pearled axon morphology in live neurons from organotypic slice cultures. Scale bar: 500 nm. Individual axons are straightened and represented on the right panels. Scale bar: 200 nm.

Extended Data Fig. 2 Griswold, et al.

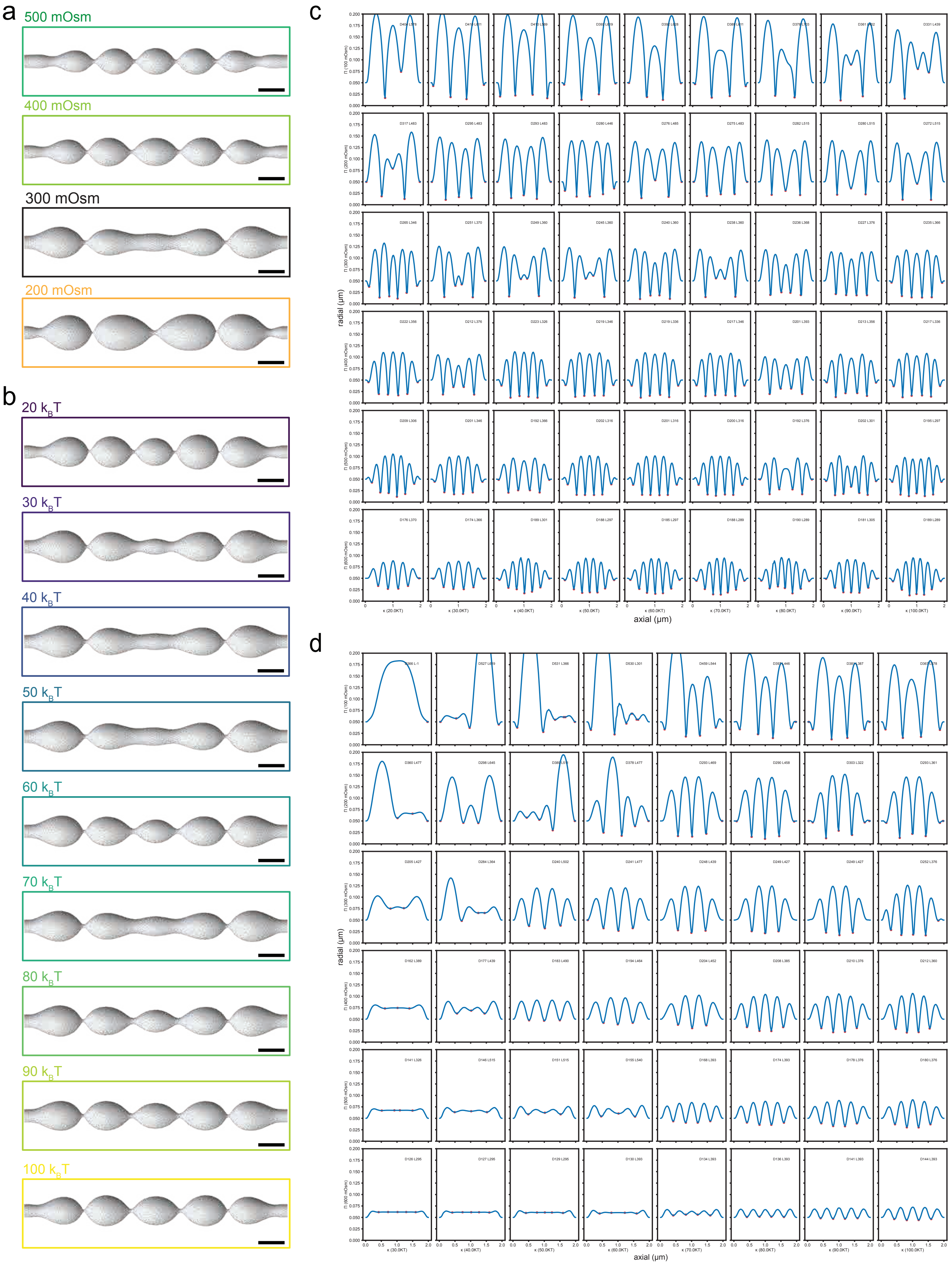

**Extended Data Fig. 2**, related to Fig. 2.

a. Axon morphology is modeled using classic Helfrich membrane model and governed by the membrane bending, surface tension, and osmotic condition. 3D model prediction of axon morphology in indicated osmotic conditions, tension of 0.001 mN/m, and bending rigidity of 50$k_B$T. Scale bar: 200 nm.

b. 3D model prediction of axon morphology at indicated bending rigidity, tension of 0.001 mN/m, and osmolarity 300 mOsm. Scale bar: 200 nm.

c. Rotationally averaged shape profiles of axons under constant 0.001 mN/m tension and varying osmotic conditions and bending rigidity. The bead diameter (D) and length (L) in nanometers are given as a text inset for each condition.

d. Rotationally averaged shape profiles of axons under constant 0.01 mN/m tension and varying osmotic conditions and bending rigidity. The bead diameter (D) and length (L) in nanometers are given as a text inset for each condition.

Extended Data Fig. 3 Griswold, et al.

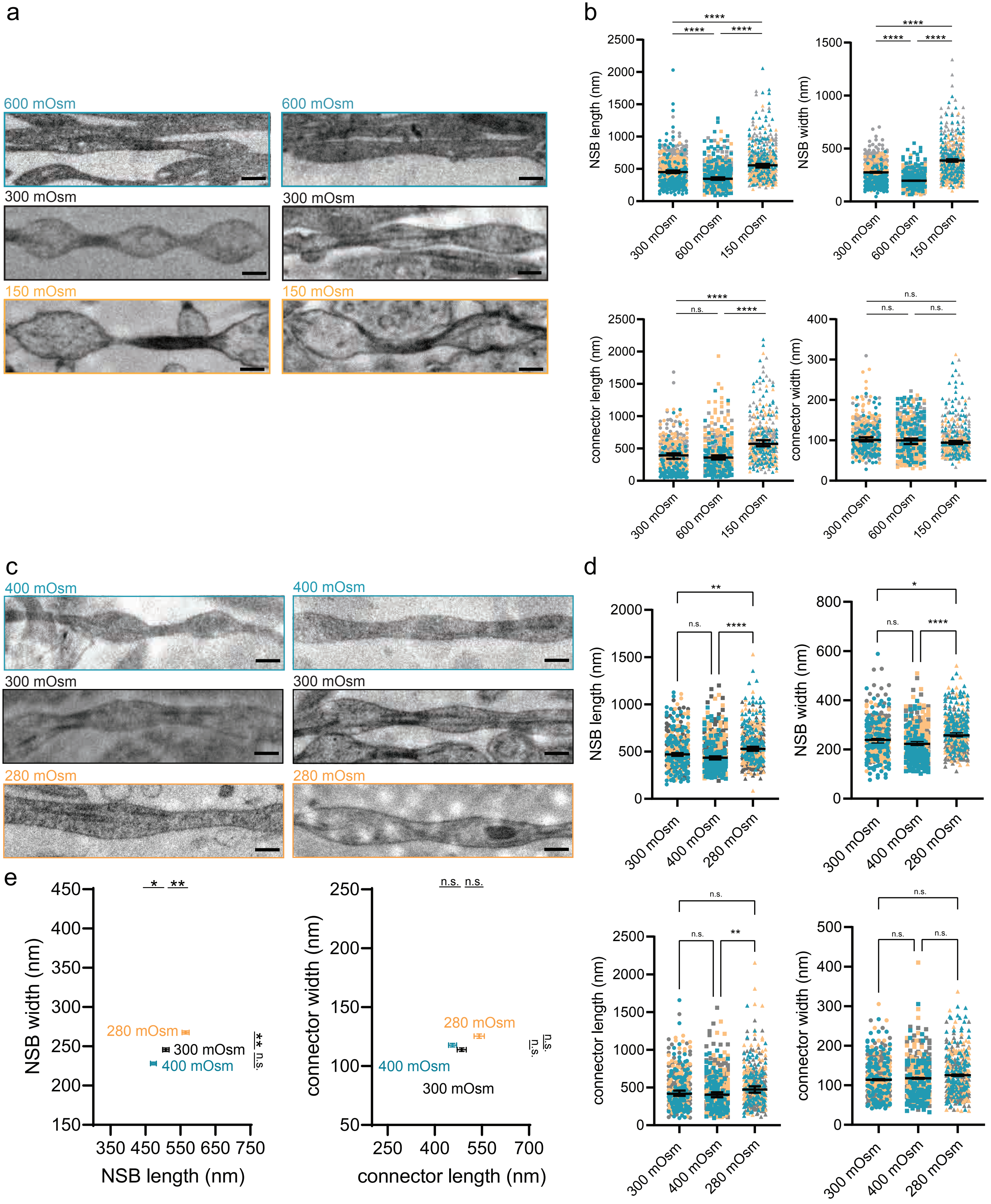

**Extended Data Fig. 3**, related to Fig. 2

a. Additional example micrographs of axons high-pressure frozen at indicated osmotic conditions. Scale bar: 200 nm.

b. Super Plots of Fig. 2f, showing experimental variability. Median and 95% confidence intervals are shown. N = 3 independent cultures and n = 100 axons each. Each color represents one replicate. Each dot is one axon.

c. Example micrographs of axons high-pressure frozen at indicated osmotic conditions. Mannitol was used to adjust the osmolarity. Scale bar: 200 nm.

d. Super Plots of Extended Data Fig. 2e, showing the dimensions of NSBs at indicated osmotic conditions. Median and 95% confidence intervals are shown. N = 3 independent cultures and n = 100 axons each. Each color represents one replicate. Each dot is one axon.

e. Plot showing the dimensions of NSBs at indicated osmotic conditions. Mean and SEM are plotted. N = 3 independent cultures, n = 100 axons each.

*n.s*. not significant, ** $p<0.01$, **** $p<0.0001$.

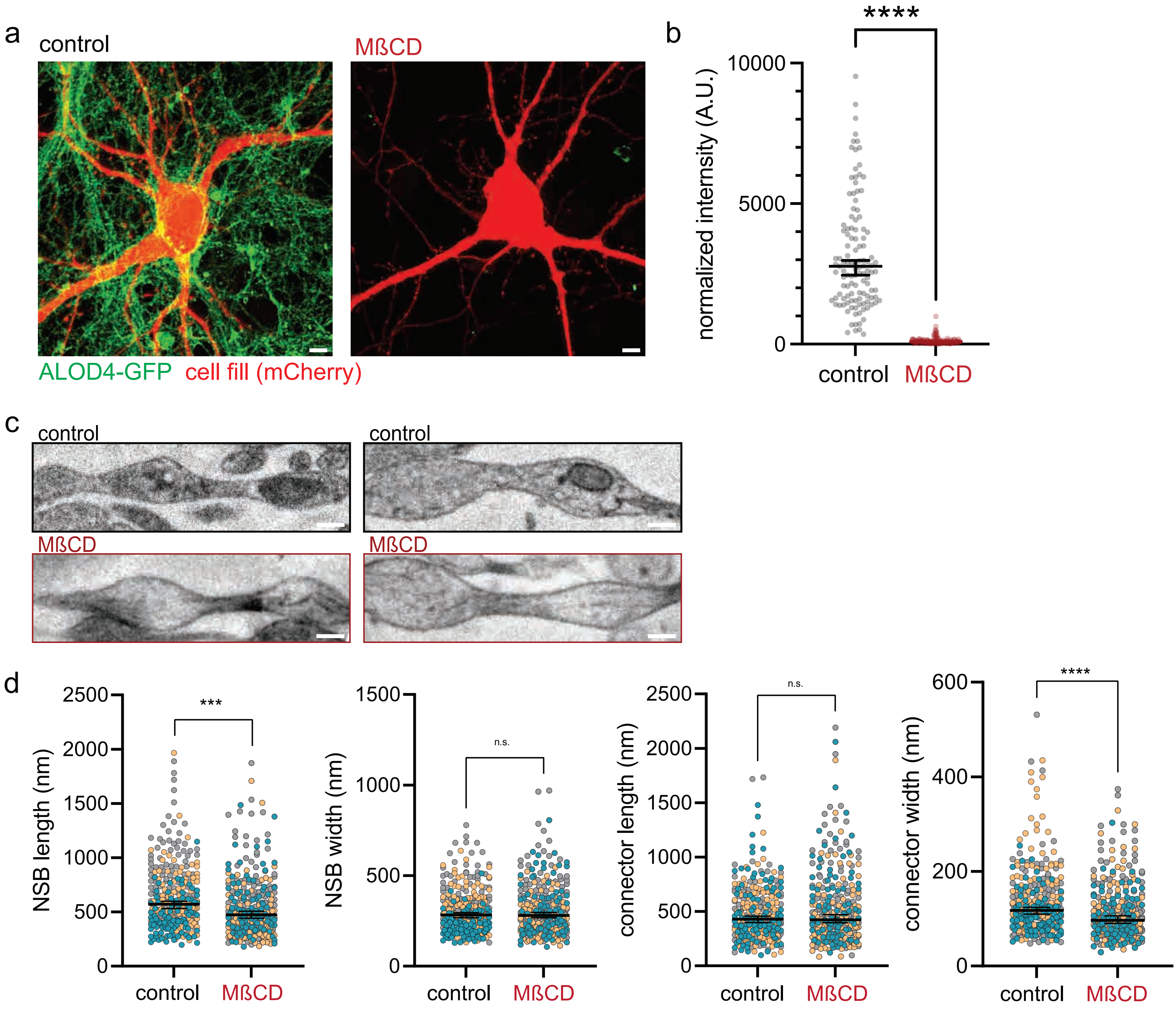
a
control
MßCD
ALOD4-GFP
cell fill (mCherry)
b
****
normalized intensity (A.U.)
10000
5000
0
control
MßCD
c
control
control
MßCD
MßCD
d
NSB length (nm)
2500
2000
1500
1000
500
0
***
control
MßCD
NSB width (nm)
1500
1000
500
0
n.s.
control
MßCD
connector length (nm)
2500
2000
1500
1000
500
0
n.s.
control
MßCD
connector width (nm)
600
400
200
0
****
control
MßCD

**Extended Data Fig. 4**. related to Fig. 2

a. NeonGreen-ALOD4 staining of cultured mouse hippocampal neurons treated with sham (control) or 5 mM MßCD for 30 min. Scale bar = 5 µm.

b. Plot showing the normalized intensity of NeonGreen-ALOD4. Signals are normalized by the area of each region-of-interest. Control: 2773 ± 234 A.U., MβCD: 86 ± 13 A.U., median and 95% confidence intervals are plotted. A.U. arbitrary unit.

c. Additional example micrographs of axons high-pressure frozen at indicated osmotic conditions. Scale bar: 200 nm.

d. Super Plots of Fig. 2h, showing variability. N = 3 independent cultures and n = 100 axons each median and 95% confidence intervals are plotted. *n.s.* not significant, ****P* < 0.001, **** *P* < 0.0001. Each color represents one replicate. Each dot is one axon.

Extended Data Fig. 5 Griswold, et al.

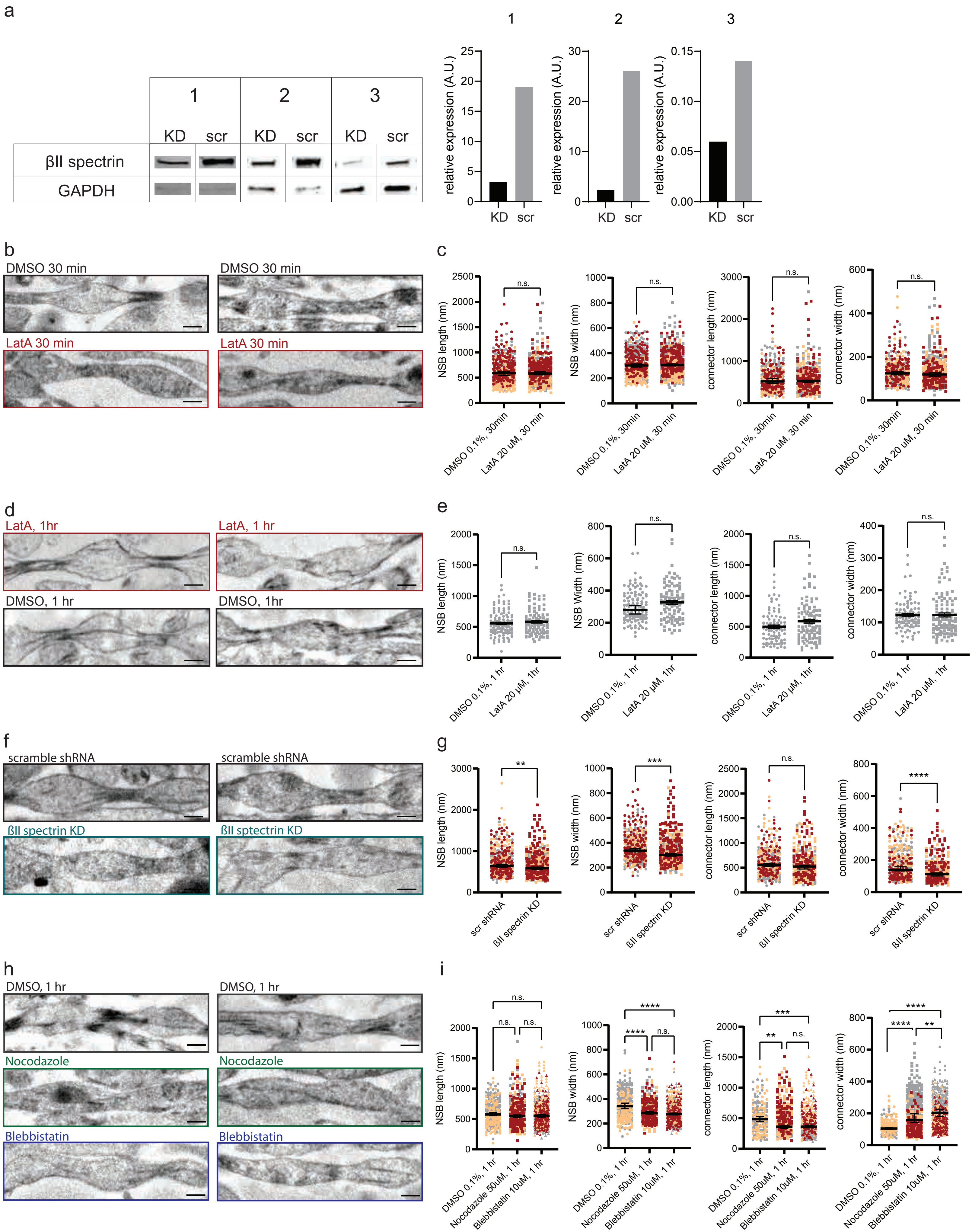

**Extended Data Fig. 5**, related to Fig. 3

a. Western blots and quantification showing the efficiency of shRNA-mediated knock down (KD) of ßII spectrin. Scr, scramble.

b. Additional example micrographs of axons from neurons treated with indicated drugs. Scale bar: 200 nm.

c. Super Plots of Fig. 3a, showing experimental variability. Median and 95% confidence intervals are shown. N = 3 independent cultures and n = 100 axons each. Each color represents one replicate. Each dot is one axon.

d. Additional example micrographs of axons from neurons treated with indicated drugs. Note that these neurons are treated with DMSO or Latrunculin A for 1 hour. Scale bar: 200 nm.

e. Super Plots, showing experimental variability of d. Median and 95% confidence intervals are shown. N = 1 and n = 100 axons. Each dot is one axon.

f. Additional example micrographs of axons from neurons infected with lentivirus carrying scramble shRNA or shRNA against ßII spectrin. Scale bar: 200 nm.

g. Super Plots of Fig. 3d, showing experimental variability. Median and 95% confidence intervals are shown. N = 3 independent cultures and n = 100 axons each. Each color represents one replicate. Each dot is one axon.

h. Additional example micrographs of axons from neurons treated with indicated drugs. Scale bar: 200 nm.

i. Super Plots of Fig. 3f, showing experimental variability. Median and 95% confidence intervals are shown. N = 3 independent cultures and n = 100 axons each.

*n.s.*, not significant, $^{**}P < 0.01$,$^{***}P < 0.001$, $^{****}P < 0.0001$.

Extended Data Fig. 6 Griswold, et al.

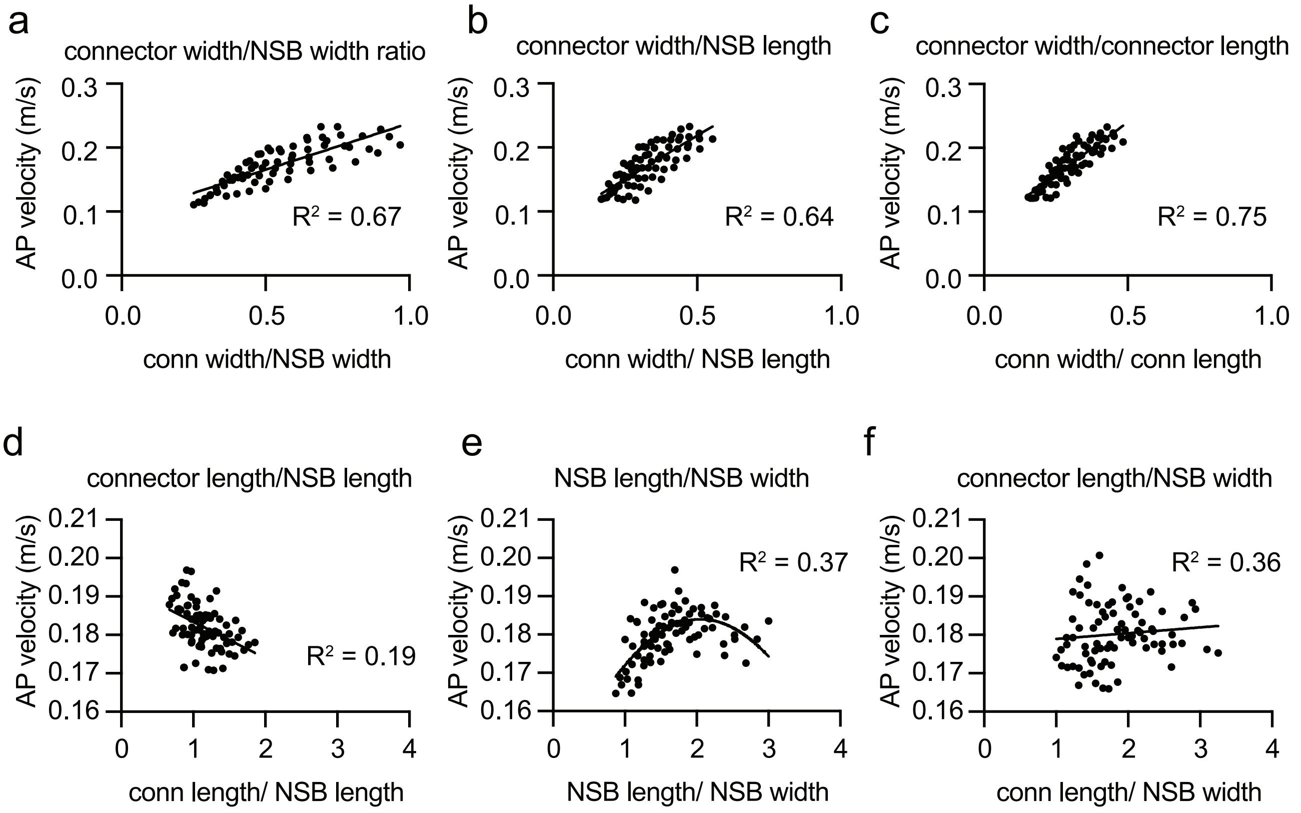

**Extended Data Fig. 6**, related to Fig. 4

a-f, Plots showing the relationship between AP velocity and the ratio of two measured values like connector width/NSB width (a), connector width/NSB length (b), connector width/connector length (c), connector length/NSB length (d), NSB length/NSB width (e), and connector length/NSB width (f). The data are fitted by a simple linear regression in all except for e, which is fitted by a gaussian non-linear curve.

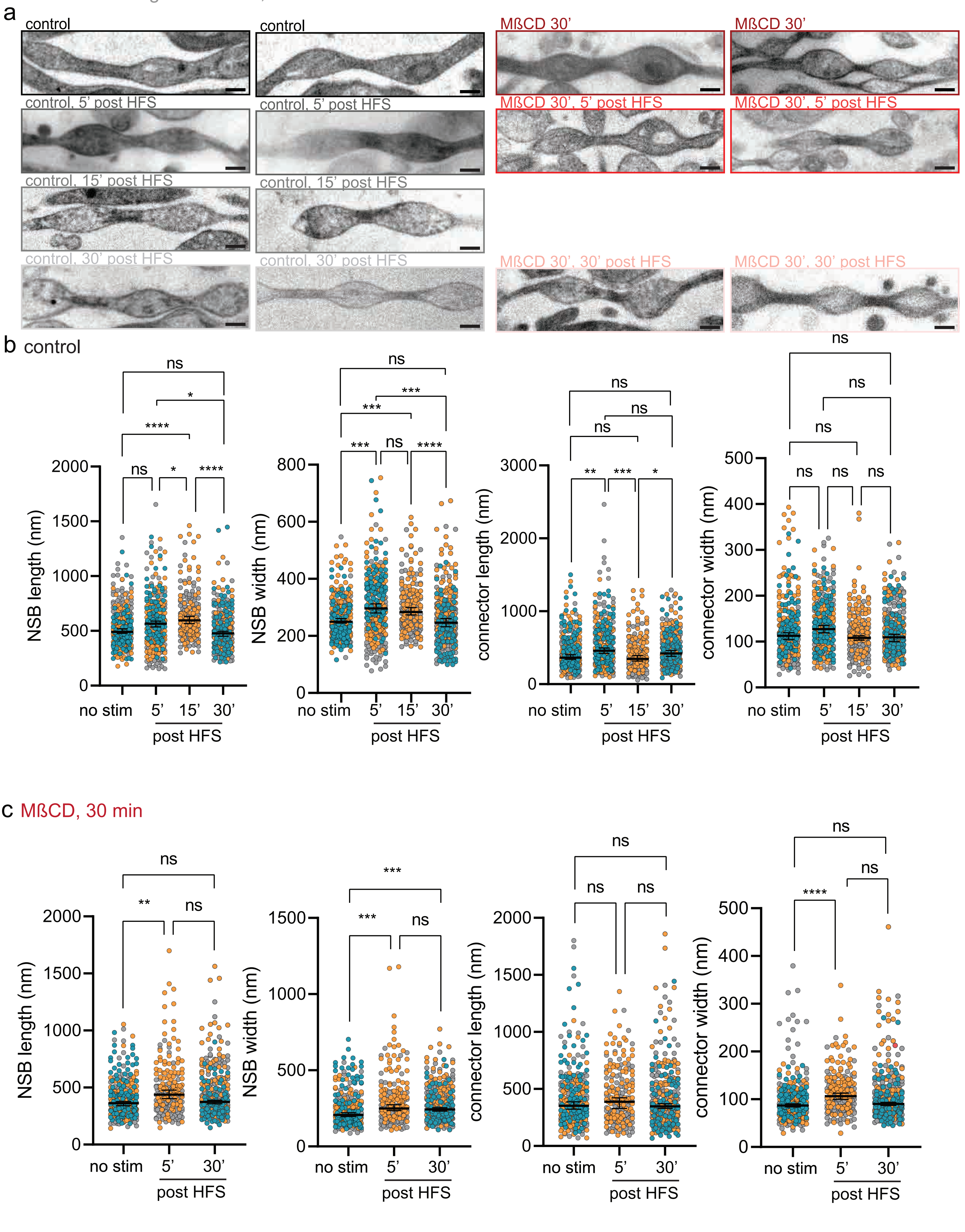

a
control
control
MßCD 30'
MßCD 30'
control, 5' post HFS
control, 5' post HFS
MßCD 30', 5' post HFS
MßCD 30', 5' post HFS
control, 15' post HFS
control, 15' post HFS
control, 30' post HFS
control, 30' post HFS
MßCD 30', 30' post HFS
MßCD 30', 30' post HFS
b control
NSB length (nm)
NSB width (nm)
connector length (nm)
connector width (nm)
no stim
5'
15'
30'
post HFS
ns
*
****
**
***
c MßCD, 30 min
NSB length (nm)
NSB width (nm)
connector length (nm)
connector width (nm)
no stim
5'
30'
post HFS

**Extended Data Fig. 7**, related to Fig.5

a. Additional example micrographs showing axon morphology from neurons unstimulated or stimulated with 3x100 pulses at 100 Hz (high-frequency stimulation, HFS) and high-pressure frozen at 5 min and 30 min after stimulation. Scale bar: 200 nm

b. Super Plots of Fig. 5b, showing experimental variability. Median and 95% confidence intervals are shown. N = 3 independent cultures except for control 15' post stim and MβCD 5' post stim which were N = 2. n = 100 axons each. Each color represents one replicate. Each dot is one axon. *n.s.*, not significant, *$P < 0.05$, **$P < 0.01$,***$P < 0.001$, ****$P < 0.0001$
