## Supplementary Info for "Membrane mechanics dictate axonal morphology and function"

We note that all codes and softwares for reproducing this work are available from `https://github.com/RangamaniLabUCSD/2023-axon-pearling` and archived at `http://doi.org/10.5281/zenodo.8060707`.

### 1 Axon pearling simulations

#### 1.1 Equilibrium conditions for tubes

Following the approach outlined by Refs. (*1*, *2*), we first establish the general conditions for pulling a tube from a reservoir of membrane. The general form of the energy of a membrane with an external force $f$ acting over length $L$ is,

$$F = \int_A \left[\kappa(H - c_0)^2 + \sigma\right] dA - pV - fL. \tag{1}$$

Where $H$ is the mean curvature give by $(c_1 + c_2)/2$ In the special case of a cylinder which has $c_1 = 1/R$ and $c_2 = 0$, assuming that $p = 0$, and $f \neq 0$,

$$F = \left[\kappa\left(\frac{1}{2R} - c_0\right)^2 + \sigma\right] 2\pi RL - fL. \tag{2}$$

At equilibrium the energies are minimized with respect to the radius and length of the tube such that

$$\frac{\partial F}{\partial R} = 0, \quad \frac{F}{\partial L} = 0. \tag{3}$$

Considering first the minimization with respect to the radius,

$$\begin{aligned} 0 = \frac{\partial F}{\partial R} &= 2\pi L\left[\frac{-\kappa}{4R^2} + \kappa c_0^2 + \sigma\right] \\ &= -\frac{\kappa}{4R^2} + \kappa c_0^2 + \sigma \end{aligned}$$

Thus,

$$R = \frac{1}{2\sqrt{c_0^2 + \sigma/\kappa}}, \tag{4}$$

relates the radius to the spontaneous curvature, tension, and bending rigidity; moreover,

$$c_0 = \sqrt{1/4R^2 - \sigma/\kappa}. \tag{5}$$

Assuming an initial geometry of a cylinder with radius $R = 0.05$ μm and given values of tension and bending rigidity, we can obtain the homeogenous spontaneous curvature to sustain the tube.

#### 1.2 Osmotic pressure relationship and computing pressure force

The osmolarity of the solution, $\bar{c}$, is given by

$$\bar{c} = \sum_i \varphi_i k_i C_i, \tag{6}$$

where $\varphi$ is the osmotic (activity) coefficient, $k$ is the number of dissociated ions, and $C$ is the concentration.

The osmotic pressure is given by

$$P = RT\bar{c} = RT \sum_i \varphi_i k_i C_i \tag{7}$$

Simplifying by homogenizing the contributions of each ionic species, the osmotic pressure energy is given by,

$$E_p = \int_{\bar{V}}^{V} \Delta P(\tilde{V}; \vec{r}) d\tilde{V} = \int_{\bar{V}}^{V} kRT \left( \frac{n}{V} - \bar{c} \right) dV = kRTn[\bar{c}V/n - \ln(\bar{c}V/n) - 1], \tag{8}$$

where $RT$ is the ideal gas constant and temperature, $n$ is the amount of enclosed solute within the geometry, $\bar{c}$ is the concentration of the external solvent, and $V$ is the current volume.

To define the initial molar amount of enclosed solute, we assume that the interior of the cell is at the conventional $c_{t=0} = 300$ mOsm. The amount of solute is given by the product of concentration and initial volume, $n = c_{t_0} V_{t_0}$. Since the initial geometry, a cylinder with radius 0.05 μm, is somewhat arbitrary, we scale the initial volume by a factor of three to set the effective target volume,

$$n = c_{t_0} \cdot 3V_{t_0}. \tag{9}$$

An effect of this scaling is that the initial volume is much different than the target volume. The contributions from high pressure can lead to numerical instabilities. We assume that since we are modeling only a small section of axon with the majority of the cell volume not represented, that the bulk of the cell also responds to osmotic pressure. To capture the contribution implicitly, we scale the energy by a reservoir volume $V_r$ nondimensionalized by a unit volume,

$$E_p = \frac{kRTn}{V_r}[\bar{c}V/n - \ln(\bar{c}V/n) - 1]. \tag{10}$$

For our intuition, we note that at nearly isoosmotic conditions,

$$\lim_{V \to \bar{V}} kRTn[\bar{c}V/n - \ln(\bar{c}V/n) - 1] \approx \frac{1}{2} K_v \frac{(V - \bar{V})^2}{\bar{V}^2} + \mathcal{O}(V^3). \tag{11}$$

we can define a constant $K_v = kRTn$ which accounts for the constant parameters. Thus the reservoir volume is serving to temper the effective bulk modulus.

#### 1.3 Parameter variation and other simulation conditions

We vary the range of osmolarities from 100–600 mOsm, bending modulus 20–100 $k_{\mathrm{b}}T$, and tension 0.001–0.01 mN m. For all simulations, we assume that the temperature is 310 K. As aforementioned, the initial condition for all conditions is a cylinder with initial radius, $R = 0.05$ μm. The spontaneous curvature is homeogeneous over all vertices with value computed by Eq. (5). We use a reference volume of 700 μm$^3$ from the literature (c.f., Larsen *et al.* (*3*)). All systems are simulated under constant tension conditions. The contribution to system energy from osmotic pressure is given by Eq. (10); Thus the system tends towards a volume which produces a condition isoosmotic with the prescribed solvent osmolarity. The relaxation of the membrane systems with given material properties and osmotic conditions were simulated using *Mem3DG* version 0.0.6rc2 (*4*). Fixed boundary conditions were used for all simulations (i.e., vertices within the first 2 rings to a boundary have fixed position). The intial timestep for each system is set relative to the given bending modulus and allowed to adaptively vary with system condition. The system geometry is propagated using forward Euler integration. To prevent mesh degradation, the mesh is conditioned every 100 timesteps. Resulting configurations are stored in intervals of 2000 timesteps, which can vary in size. Simulations are terminated by one of three conditions: 1) reaching a L2 force threshold tolerance of 1 × 10$^{-9}$ pN$^2$, 2) reaching 1e7 timesteps, or 3) the adaptive timestep becomes too small.

##### 1.4 Analysis of the output geometries

2D profiles of the output geometries were generated by rotationally averaging the output vertex positions. The maximum value of each profile is used to obtain the diameter of the NSB. The length of each NSB is computed by the mean distance between local minima of each profile. All profiles are shown in Supplemental Figure 2.

#### 2 Action potential propagation

To model the action potential propagation in the axon, we follow the work of Hodgkin and Huxley (*5*) and use the cable equation given by

$$i = \frac{1}{\rho}\frac{\partial^2 V}{\partial x^2}, \tag{12}$$

where $i$ is the membrane current per unit length, $V$ is the displacement of the membrane potential from its resting value, $x$ is the distance along the axon, and $\rho$ is the internal resistance per unit length. To transform the unit length to density, we take $i = 2\pi\Delta x r I$ and $\rho = R\pi r^2/\Delta x$, where $\Delta x$ is the unit length, $r$ the axon radius, $R$ the specific resistance of the axoplasm, and $I$ the total membrane current density. Then Eq. 12 can be rewritten as

$$I = \frac{r}{2R}\frac{\partial^2 V}{\partial x^2}. \tag{13}$$

The total current can be divided into a capacity current and an ionic current $I_i$, hence

$$I = C_M\frac{\mathrm{d}V}{\mathrm{d}t} + I_i = C_M\frac{\mathrm{d}V}{\mathrm{d}t} + I_{Na} + I_K + I_l. \tag{14}$$

Here, $C_M$ is the membrane capacity per unit area and the ionic current is split into components carried by sodium ($I_{Na}$), potassium ($I_K$), or other ions ($I_l$). Using Ohm's law, these currents can be linearly related to the driving potential:

$$I_j = G_j(V(t),t)(V(t) - E_j), \quad j \in \{Na, K, l\}, \tag{15}$$

where $E_j$ is the reversal potential and $G_j$ the conductance. For the leak current $I_l = G_m(V - E_l)$, with $G_m$ constant. For the sodium and potassium currents, the ionic conductances are expressed as the maximum conductance, $G_j$, multiplied by a number that represents the open fraction of $G_j$. Such a number is obtained from the dynamics of activating and inactivating gating particles (*5*, *6*). For the potassium current $I_K = G_K n^4(V - E_K)$, with $n$ representing an activating particle that has the following dynamics:

$$\frac{\mathrm{d}n}{\mathrm{d}t} = \alpha_n(V)(1-n) - \beta_n(V)n, \quad \alpha_n = \frac{10-V}{100(\mathrm{e}^{(10-V)/10}-1)}, \quad \beta_n = 0.125\mathrm{e}^{-V/80}. \tag{16}$$

The sodium current $I_{Na} = G_{Na} m^3 h(V - E_{Na})$ has an activation and inactivation particle, $m$ and $h$, respectively, described by

$$\frac{\mathrm{d}m}{\mathrm{d}t} = \alpha_m(V)(1-m) - \beta_m(V)m, \quad \alpha_m = \frac{25-V}{10(\mathrm{e}^{(25-V)/10}-1)}, \quad \beta_m = 4\mathrm{e}^{-V/18}, \quad \text{and} \tag{17}$$

$$\frac{\mathrm{d}h}{\mathrm{d}t} = \alpha_h(V)(1-h) - \beta_h(V)h, \quad \alpha_h = 0.07\mathrm{e}^{-V/20}, \quad \beta_h = \frac{1}{\mathrm{e}^{(30-V)/10}-1}. \tag{18}$$

We simulate the cable equation (Eq. (12)) with the currents described by Hodgkin and Huxley for 10 ms in COMSOL Multiphysics 6.1. The parameter values are taken from (*6*) for the model and from experiments for the axon shape, see Table 1. The first 90 µm segment of the axon corresponds to the initial segment where there is no beading. After the initial segment, the axon geometry is composed of ellipsoids, representing non-synaptic boutons (NSB), connected by cylinders, representing connectors. An external current $I_{ext}$ is applied after 0.5 ms for 1 ms to the first 40 µm of the initial segment. We measured $V$ at discrete locations in the axon throughout the simulation. To calculate the action potential velocity we obtained the time when $V$ has its maximum value at the different locations and divide the distance of the measuring locations by the difference between maximum times. The simulations have the mean geometrical parameters, unless stated otherwise: NSB length = 475 nm, NSB width = 300 nm, connector length = 525 nm, and connector width = 162.5 nm.

The periodic location of sodium channels is modeled by multiplying the potassium activation variable times $f(x) = -0.5\mathrm{sign}(\sin(2\pi x/190) - 0.4) + 0.5$ ($f(x) = 0.5\mathrm{sign}(\sin(2\pi x/190) - 0.3) + 0.5$). This results in a periodic (190 nm) step-like function that equals 1 for $\sim 2/3$ of the period, which resembles experimental data (*7*). To conserve the amount of sodium in the channels, the conductance was scaled $G_{Na} = G_{Na}(1 + l_{off}/l_{on})$, where $l_{off}$ ($l_{on}$) is the percentage of surface area without (with) sodium channels.

**Table 1: Model Parameters**

| Symbol | Definition | Value |
|---|---|---|
| $C_M$ | membrane capacity | 1 μF cm$^{-2}$ |
| $R$ | specific resistance of the axoplasm | 70.8 Ω cm |
| $I_{ext}$ | external current | 30 μA cm$^{-2}$ |
| $G_K$ | maximum potassium conductance | 36 mS cm$^{-2}$ |
| $G_{Na}$ | maximum sodium conductance | 120 mS cm$^{-2}$ |
| $G_m$ | leak conductance | 0.3 mS cm$^{-2}$ |
| $E_K$ | ionic reversal of potassium | −12 mV |
| $E_{Na}$ | ionic reversal of sodium | 115 mV |
| $E_l$ | ionic reversal of leak | 10.613 mV |
| $V_0$ | initial displacement of the membrane potential from its resting value | 0 mV |
| $n_0$ | initial value of potassium activation | 0.32 |
| $m_0$ | initial value of sodium activation | 0.05 |

### References


1. I. Derényi, F. Jülicher, J. Prost, *Physical Review Letters* **88**, 238101, DOI `10.1103/PhysRevLett.88.238101` (May 28, 2002).
2. C. R. Shurer *et al.*, *Cell* **177**, 1757–1770.e21, ISSN: 0092-8674, DOI `10/ggctwr` (June 13, 2019).
3. N. Y. Larsen *et al.*, *Communications Biology* **4**, 1–15, ISSN: 2399-3642, DOI `10.1038/s42003-021-02548-6` (1 Sept. 2, 2021).
4. C. Zhu, C. T. Lee, P. Rangamani, *Biophysical Reports* **2**, 100062, ISSN: 26670747, DOI `10.1016/j.bpr.2022.100062`, pmid: `36157269` (Sept. 14, 2022).
5. A. L. Hodgkin, A. F. Huxley, *The Journal of Physiology* **117**, 500 (1952).
6. C. Koch, *Biophysics of computation: information processing in single neurons* (Oxford university press, 2004).
7. K. Xu, G. Zhong, X. Zhuang, *Science* **339**, 452–456 (2013).
